# Microbial metabolism of food allergens determines the severity of IgE-mediated anaphylaxis

**DOI:** 10.1101/2025.02.17.638013

**Authors:** Elisa Sánchez-Martínez, Liam E. Rondeau, Manuel Garrido-Romero, Bruna Barbosa da Luz, Dominic A. Haas, Gavin Yuen, Dana Coppens, Peter Hall, Rebecca Dang, Xuan-Yu Wang, Lucía Moreno-Serna, Celia López-Sanz, Emilio Nuñez-Borque, Maria Garrido-Arandia, Araceli Diaz-Perales, Yolanda R. Carrasco, Joshua F.E. Koenig, Tina D. Walker, Manel Jordana, Elena F. Verdu, Premysl Bercik, Giada De Palma, Michael G. Surette, Pedro Ojeda, Francisco Vega, Carlos Blanco, Lingdi Zhang, Supinda Bunyavanich, Wayne G. Shreffler, Sarita U. Patil, F. Javier Moreno, Rodrigo Jiménez-Saiz, Alberto Caminero

## Abstract

Anaphylaxis is an acute, potentially life-threatening reaction, often triggered by foods and largely mediated by IgE. Critically important to anaphylaxis are the factors that modulate its severity. The human microbiota is known to influence oral tolerance, but the microbial mechanisms directly involved in IgE-mediated anaphylaxis remain unknown. Here, we demonstrate that human saliva and jejunum harbor peanut-degrading bacteria that metabolize immunodominant allergens (Ara h 1 and 2) and alter IgE-binding. Additionally, we provide *in vivo* evidence that oral bacteria metabolize peanut allergens, influencing systemic allergen exposure and anaphylaxis severity. Finally, in clinical studies, we observe that common peanut-degrading bacteria, such as *Rothia,* from the oral cavity, are more abundant in peanut-allergic patients who exhibit better tolerance to allergen exposure. Altogether, these results demonstrate that human microbiota modulates IgE-mediated reactions through allergen metabolism. We reveal a novel microbial mechanism with potential to prevent, or reduce, the severity of IgE-mediated anaphylaxis.

## INTRODUCTION

Inflammation typically restores homeostasis in response to tissue damage or infection but can cause immunopathology when dysregulated^1^. Anaphylaxis exemplifies this, with acute and potentially fatal outcomes within minutes. The main pathway of anaphylaxis is mediated by immunoglobulin (Ig)E and mast cells: upon allergen binding, mast cell-bound IgE induces the rapid release of inflammatory mediators like tryptase or leukotrienes, which drive the acute clinical manifestations of anaphylaxis^2-4^. Foods are common anaphylaxis triggers, with peanut (PN) being a leading cause of food-induced anaphylaxis and allergy-related deaths among children^5,6^. Food-anaphylaxis, particularly PN-triggered, is most common in North America and Europe and less frequent in the Asia-Pacific^7,8^. The burden of PN allergy is aggravated by its persistence in >70% of individuals, the absence of curative treatments, and the frequent accidental exposures despite patients’ avoidance efforts^9-12^.

The severity of food-induced anaphylaxis is not solely determined by allergen-specific IgE levels, as clinical reactivity is influenced by a complex interplay of genetic, environmental, dietary, and behavioral factors, as well as co-factors and comorbidities^13,14^. Notably, clinical observations often reveal a disconnect between serum levels of allergen-specific IgE—the key molecule involved in food-induced anaphylaxis—and clinical reactivity^15-17^, underscoring the importance of elucidating additional modulators of anaphylaxis^2^. The human microbiota has gained considerable attention in this regard for its capacity to influence both oral tolerance and dietary antigen immunogenicity^18-22^. Indeed, studies have revealed differences in the intestinal microbiota composition of food-allergic patients^23-25^. Nonetheless, the microbial mechanisms involved in food-induced anaphylaxis remain largely unknown.

The oro-gastrointestinal microbiota is often regarded as a “second metabolic organ” capable of breaking down dietary components that are otherwise resistant to human digestive enzymes. Certain food allergens, including the immunodominant PN allergens Ara h 1 and 2^26,27^, resist complete digestion by mammalian digestive enzymes^28,29^. Here, we demonstrate that human saliva and the small intestine contain PN-degrading bacteria capable of metabolizing immunodominant allergens and modulating IgE-specific immunity. We also show the *in vivo* capacity of these bacteria to participate in PN-metabolism, thereby modulating systemic allergen access and IgE-mediated anaphylaxis. Finally, in two clinical studies, we describe that common PN-degrading bacteria, such as *Rothia* from the oral cavity, are more abundant in allergic patients who exhibit higher allergen thresholds to controlled allergen exposure. These results demonstrate the human microbiota’s ability to modulate IgE-mediated reactions through allergen metabolism and reveal a novel microbial mechanism with translational potential to prevent or mitigate IgE-mediated anaphylaxis.

## RESULTS

### Microbiota participates in peanut allergen metabolism

To study the role of microbes in PN metabolism, we used C57BL/6 mice with distinct microbiota compositions: germ-free (GF), specific pathogen-free (SPF; diverse), and minimal microbiota (MM; derived from Altered-Schaedler Flora, stable with limited species) (**Figure S1A-C**). Mice were gavaged with PN and sacrificed after 40 minutes (**Figure 1A**). The quantities of PN allergens Ara h 1 and 2 were higher in small intestinal content of GF and MM mice, compared to mice with diverse SPF microbiota (**Figure 1B**). Moreover, microbial alpha-diversity across groups correlated inversely with Ara h 1 and 2 in the small intestine (**Figure S1D**). We also detected variations in the concentrations of these allergens systemically. Compared to SPF mice, GF and MM mice presented higher serum levels of Ara h 1 but lower serum levels of Ara h 2 (**Figure 1C**). To determine whether these differences were due to variations in the digestive capacity, intestinal contents of PN-naïve mice were incubated with PN (crude PN extract, CPE) *ex vivo*. The intestinal contents of SPF mice degraded Ara h 1 and 2 more effectively than those of GF and MM mice (**Figure 1D**). We hypothesized that microbial metabolism influenced the differences observed in PN allergen digestion and attempted to isolate PN-degrading bacteria from SPF and MM mouse intestinal contents. Notably, PN-degrading bacteria were present in the small and large intestines of SPF mice but not found in MM mice. While various PN-degrading bacterial species were identified, their capacity to digest Ara h 1 and 2 varied by bacterial strain (**Figure 1E**). Together, these results demonstrate that the intestinal microbiota metabolizes PN allergens and influences their systemic availability.

**Figure 1.**
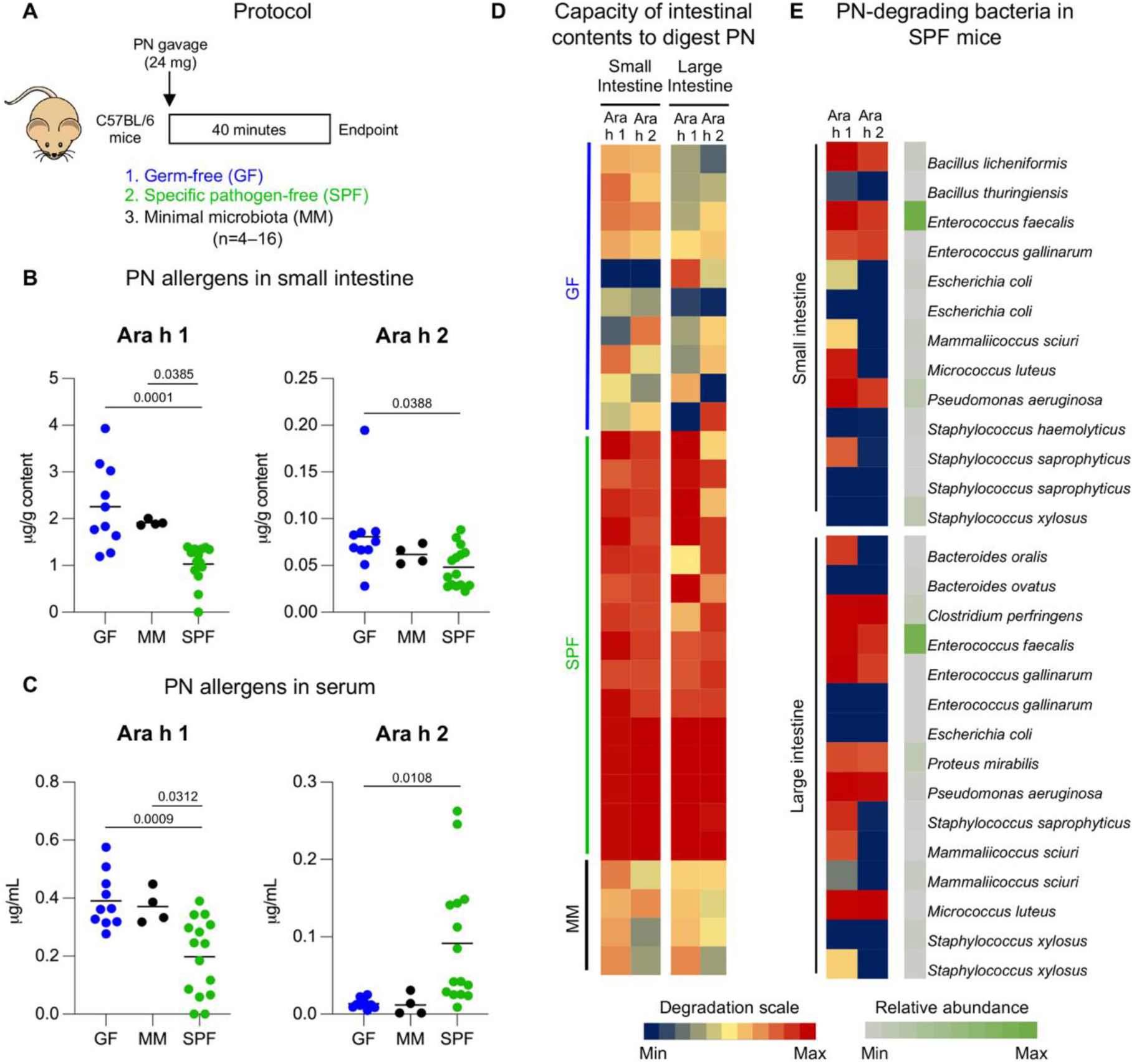
Microbiota participates in allergen metabolism *in vivo*. (A) Experimental design. Germ-free (GF), specific pathogen-free (SPF), and minimal microbiota (MM) C57BL/6 mice received peanut (PN) intragastrically (i.g.). Peripheral blood and intestinal contents were collected 40 min post-PN delivery. n=4–16 mice/group; pooled from 2 independent experiments. (B–C) PN allergens (Ara h 1 and 2) in small intestinal content (B) and serum from peripheral blood (C). Each dot = one mouse. *P* values were calculated using one-way ANOVA with Tukey’s post-hoc test. (D) Heatmap showing digestion capacity of GF, MM, and SPF microbiota against Ara h 1 and 2. PN-naïve intestinal contents were incubated with PN allergens (crude PN extract, CPE) *in vitro* and remaining allergens quantified. Each row = one mouse. Degradation capacity is expressed as % degraded relative to control incubations with CPE alone (blue=0%, red=100%). Significance levels: Small intestine Ara h 1: GF vs. SPF (*P*<0.0001), MM vs. SPF (*P*=0.0290). Small intestine Ara h 2: GF vs. SPF (*P*<0.0001), MM vs. SPF (*P*<0.0001). Large intestine Ara h 1: GF vs. SPF (*P*<0.0001), MM vs. SPF (*P*=0.0289). Large intestine Ara h 2: GF vs. SPF (*P*<0.0001), MM vs. SPF (*P*<0.0024). *P* values were calculates using one-way ANOVA with Tukey’s post-hoc test. (E) Heatmap of PN-degrading isolates and abundance of bacterial isolates from the small and large intestine of SPF mice. Isolates were obtained by plating SPF intestinal contents on PN-agar and selecting colonies with hydrolytic halos, indicative of PN degradation. Isolates were incubated in liquid media with CPE to assess Ara h 1 and 2 degradation. Each row = one isolate. Allergen degradation is shown using the same blue-red scale as in (D). Relative abundance (green scale) reflects how often each species was isolated out of all PN-degrading isolates, with darker green indicating more frequently isolated species (light=low, dark=high). See also Figure S1.

### Mice with limited peanut-degrading bacteria exhibit stronger mucosal anaphylaxis markers

To investigate the impact of microbial PN metabolism on acute allergic reactions, we used MM and SPF C3H/HeN mice, a strain susceptible to type 2 immunity and anaphylaxis upon oral PN-challenge^30,31^, as C57BL/6 mice are largely resistant to oral allergen-induced anaphylaxis^3,4^. Consistent with our previous findings in C57BL/6 mice, MM mice exhibited a reduced capacity to metabolize PN compared to SPF C3H/HeN mice (**Figure S2**). MM and SPF C3H/HeN mice were sensitized to PN with cholera toxin (CT) via intragastric (i.g.) administration once per week for 6 weeks, followed by an i.g. PN-challenge to assess anaphylactic reactions to PN allergens (**Figure 2A**). MM mice, which present reduced PN-degrading bacteria, had higher serum levels of PN-specific IgE than SPF mice (**Figure 2B**). They also exhibited higher serum levels of mucosal mast cell protease-1 (mMCP-1), indicating local activation of effector cells in the gastrointestinal tract, and greater systemic hypothermia following PN-challenge (**Figure 2C–D**). To bypass the influence of microbial PN metabolism on IgE production, we sensitized SPF and MM mice to PN with CT via intraperitoneal (i.p.) injection (**Figure 2A**). PN-specific IgE and IgG1 levels of allergic donor serum used for passive sensitization were characterized (Figure **S3A**). Following passive sensitization, both groups exhibited similar serum levels of PN-specific IgE (**Figure 2E**), yet MM mice had higher serum mMCP-1 levels and developed greater hypothermia following PN-challenge (**Figure 2F–G)**. We further bypassed the endogenous capacity to generate allergen-specific IgE in the sensitization process by passively transferring serum from PN-allergic mice into SPF and MM mice (**Figure 2A**). Both mouse strains presented similar levels of PN-specific IgE in the serum after passive sensitization (**Figure 2H**). However, MM mice again exhibited higher serum levels of mMCP-1, hypothermia, and platelet-activating factor acetylhydrolase (PAF-AH) (**Figure 2I–J & Figure S3B**). These results demonstrate that the presence of certain microbes at the site of allergen challenge can protect from anaphylaxis.

**Figure 2.**
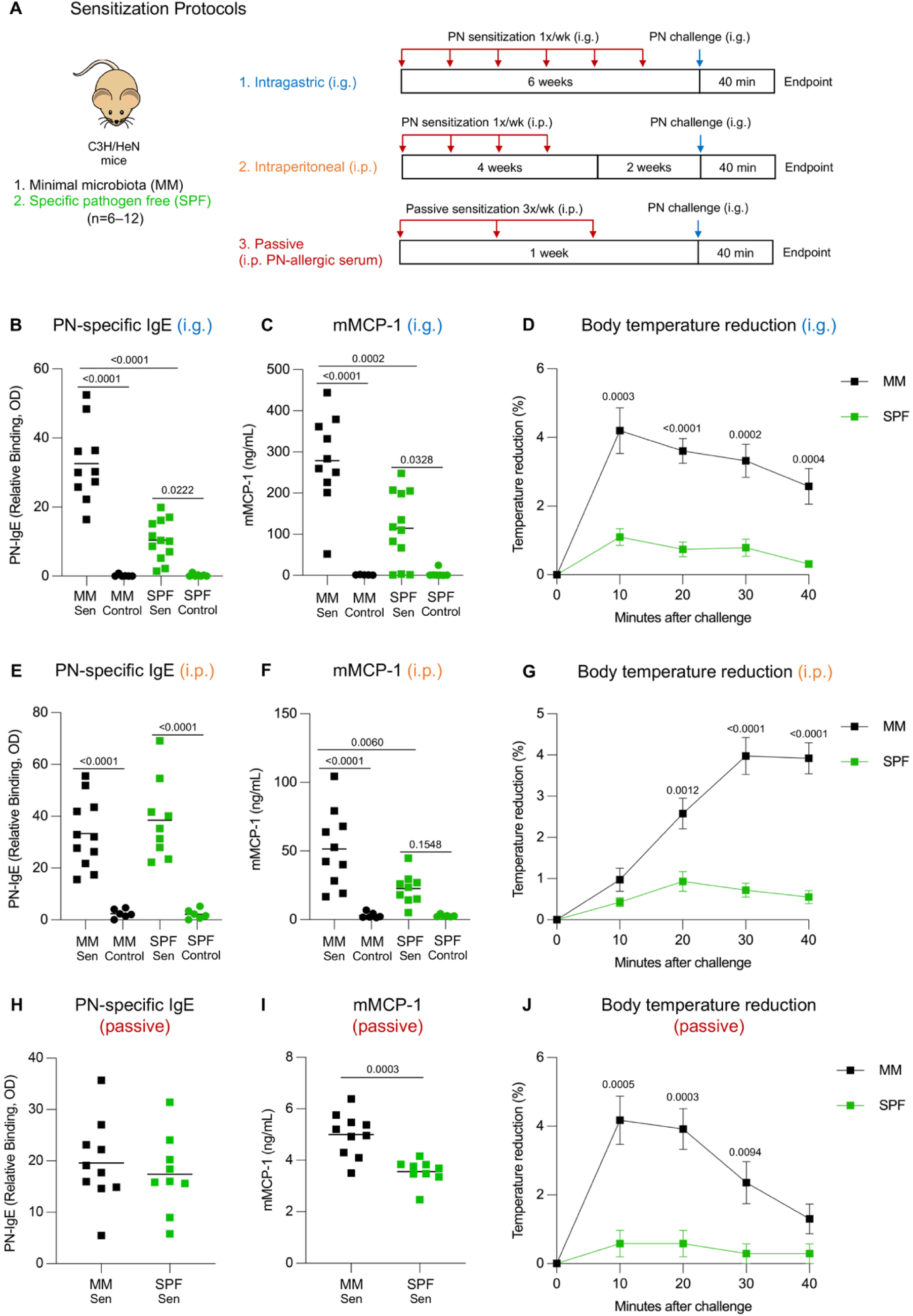
Limited microbial peanut metabolism amplifies mucosal anaphylaxis markers. (A) Experimental design for peanut (PN) sensitization and challenge of C3H/HeN mice with minimal microbiota (MM) and specific pathogen-free (SPF) microbiota. Mice were sensitized (1) intragastrically (i.g.; PN + cholera toxin) once per week for 6 weeks, (2) intraperitoneally (i.p.; PN + cholera toxin) once per week for 4 weeks, and (3) passively (i.p.; serum from PN-allergic mice) once every 48 hours, 3 times. Sensitized mice were challenged with a PN bolus i.g. and sacrificed 40 min later. (B–D) Serum PN-specific IgE (B), serum mucosal mast cell protease 1 (mMCP-1) (C), and core body temperature reduction (D) of i.g. sensitized mice after PN-challenge. (E–G) Serum PN-specific IgE (E), serum mMCP-1 (F), and core body temperature reduction (G) of i.p. sensitized mice after PN-challenge. (H–J) Serum PN-specific IgE (H), serum mMCP-1 (I), and core body temperature reduction (J) of passively sensitized mice after PN-challenge. n=9–12 mice/group for sensitized groups and n=6–7 mice/group for non-sensitized groups; pooled from 2–3 independent experiments for each comparison. Data are presented as mean where each dot = one mouse (B, C, E, F, H, I) and mean ± SEM (D, G, J). *P* values were calculated using one-way ANOVA with Tukey’s post-hoc test (B, C, E, F) and Student’s *t*-test (D, G–J). See also Figures S2–S4.

To further investigate how microbiota composition influences anaphylactic outcomes, we characterized bacterial community structure and diversity in MM and SPF mice across all sensitization protocols (**Figure S4A**). As expected, MM mice exhibited lower alpha-diversity compared to SPF mice, and this reduction in observed species was associated with increased intestinal levels of Ara h 1 and 2 following PN-challenge (**Figure S4B**). Moreover, Ara h 1 and 2 levels inversely correlate with alpha-diversity across sensitization methods (**Figure S4C**), and serum mMCP-1 concentrations showed similar negative correlations with alpha-diversity in i.g., i.p., and passive models (**Figure S4D**). These findings demonstrate that microbial complexity is linked to enhanced PN degradation and reduced effector responses.

### Human oral cavity and small intestine harbor bacteria with the capacity to digest peanut

Food-induced allergic reactions can occur within minutes of allergen exposure. Given this rapid onset, we investigated whether humans harbor PN-degrading bacteria in the oral cavity and the small intestine. We analyzed saliva from 13 volunteers and jejunum aspirates from 5 volunteers with no reported food allergies (**Table S1**). Analysis of the oral and small intestinal microbiota composition via 16S rRNA sequencing revealed donor-specific profiles, with *Prevotella*, *Rothia,* and *Fusobacterium* as the most abundant genera in the saliva and *Streptococcus* as the dominant genus in the jejunum (**Figure 3A**). Among these taxa, *Rothia* was particularly notable, accounting for up to 43% of the salivary microbiota and as much as 14% of the small intestinal microbiota. We then plated saliva samples on PN-enriched media agar to isolate bacteria with PN-degrading capacity, selecting strains that produced a hydrolytic halo, indicating their ability to degrade PN (**Figure 3B**). PN-degrading bacteria were detected in nearly all donors, with *Rothia, Staphylococcus, Streptococcus,* and *Veillonella* being the most frequently isolated genera. Each isolated bacterial strain was then incubated with CPE to facilitate allergen digestion, and the remaining Ara h 1 and 2 were quantified. Additionally, we screened a well-characterized bacterial collection for PN-degrading potential^32^. While many bacterial strains exhibited PN digestion capabilities on agar plates, their capacity to degrade PN allergens varied by taxon and strain (**Figure 3C**). *Rothia*, a dominant oral genus, consistently degraded Ara h 1 and 2 *in vitro*. This activity was observed in all tested *Rothia* strains and a phylogenetically related genus, *Micrococcus*, suggesting a conserved PN-degrading function (**Figure 3C**). In contrast, most strains from other PN-degrading genera such as *Gemella* or *Streptococcus* lacked significant allergen-degrading capacity (**Figure S5A**). While *Staphylococcus* strains showed strong PN-degrading potential, their ability to degrade PN allergens was strain-specific (**Figure 3C**). Like the oral cavity, we identified strains of *Streptococcus*, *Rothia* and *Staphylococcus* in the small intestine of healthy individuals with PN allergen-degrading capacity (**Figure S5C**). Altogether, these findings demonstrate that oral and intestinal bacteria degrade PN allergens with genus- and strain-dependent efficiency.

**Figure 3.**
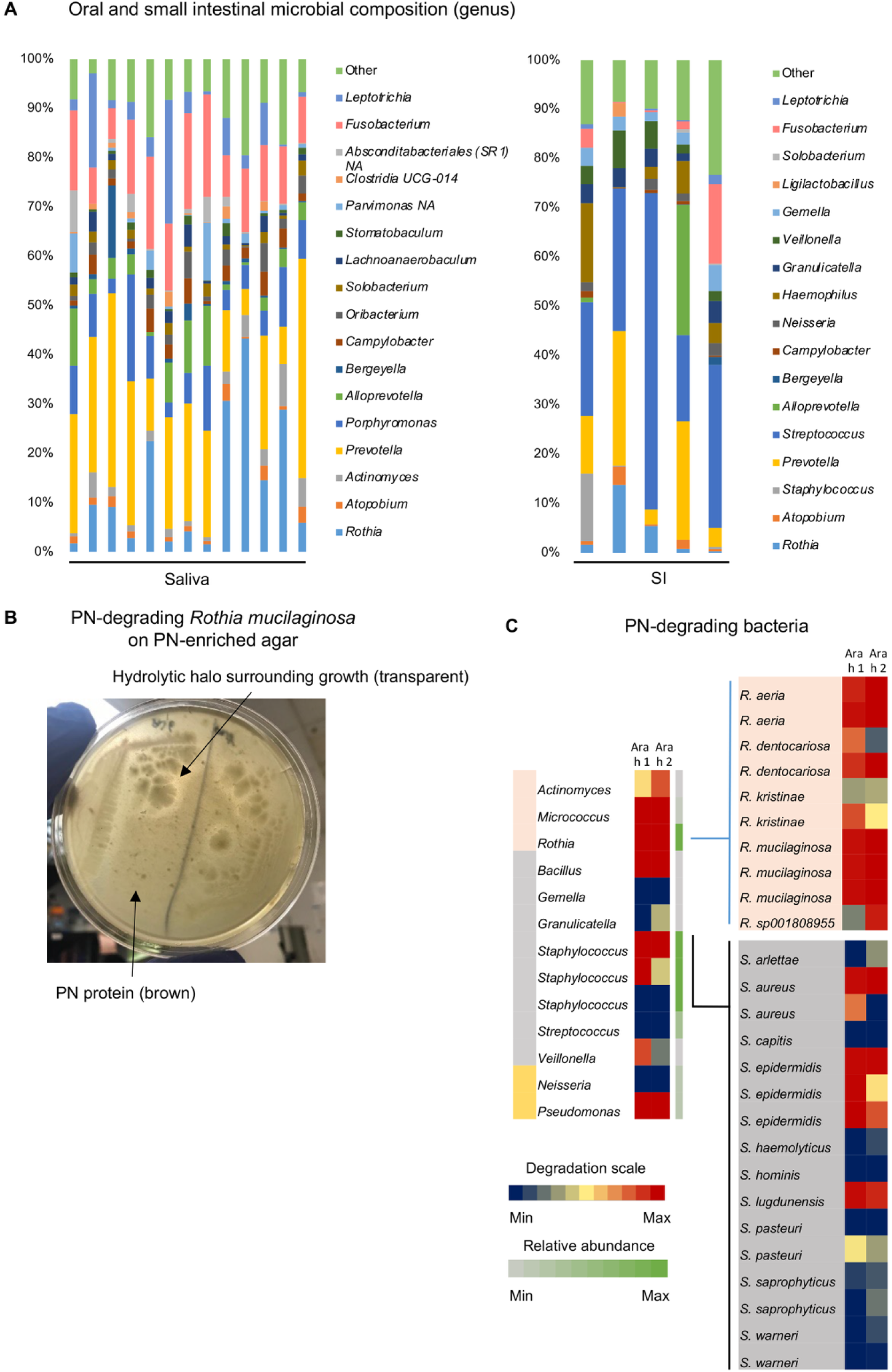
Human saliva harbors peanut-degrading bacteria. (A) Genus-level relative abundance of bacteria in saliva (n=13) and jejunal aspirates (n=5) from non-peanut (PN)-allergic individuals. Taxa <1% relative abundance not shown. (B) PN-degrading bacteria (*Rothia mucilaginosa*) from human saliva on PN-agar showing zone of clearance. (C) PN-degrading bacteria isolated from human saliva, identified for whole PN-degradation by growth on PN-agar and assessed for Ara h 1 and 2 degradation. Heatmaps show allergen degradation (blue=0%, red=100% relative to crude peanut extract control) and isolate relative abundance (green scale reflecting the frequency of isolation among all PN-degrading isolates, light=lowest, dark=highest). Data presented at the genus and species levels for *Actinomycetota* (pink), *Bacillota* (grey) and *Pseudomonota* (yellow). See also Figure S5.

### Peanut allergens are degraded by bacteria

To further study the microbial capacity for PN allergen digestion, we conducted proteomic analyses focusing on key bacterial taxa. We selected two primary genera: *Rothia*, a dominant oral taxon with efficient allergen-digestion capacity, and *Staphylococcus,* which has been previously associated with allergic diseases (**Figure S5B**)^33^. First, we confirmed the capacity of these taxa to degrade the main PN allergens, Ara h 1 (64 kDa) and Ara h 2 (17 kDa), using SDS-PAGE (**Figure 4A–B**). To strengthen these data, we performed tandem mass spectrometry analysis on the fraction below 3 kDa, screening for PN-derived peptides after bacterial digestion. Given the low relative abundance of Ara h 2 in the PN matrix (5.9-9.3% of total protein) and the noise introduced by bacterial proteins in the samples, we centered the proteomic analyses on Ara h 1 (12-16% of total protein)^34^. We identified several Ara h 1-derived peptides following PN digestion by strains of *Rothia* and *Staphylococcus*. As expected, we quantified more peptides from bacterial strains that degraded PN more extensively (*Rothia* R3, 9 peptides; S1, 11 peptides; S3, 13 peptides), according to the electrophoretic analysis. Using molecular modelling, we visualized these peptides, marking in yellow peptides digested by bacteria. (**Figure 4C–F**). Interestingly, the identified peptides shown in purple have been reported as clinically relevant IgE epitopes recognized by PN-allergic patients^35-46^. These epitopes are distributed across various regions of the 3D structure of Ara h 1, highlighting the lack of localization of the identified peptides to a single structural domain (**Figure 4C–F**). Considering the ability of these *Rothia* and *Staphylococcus* species to degrade PN, we evaluated their impact on human IgE-binding to PN after bacterial digestion by Western blotting. We used a pool of sera from PN-allergic patients with elevated levels of Ara h 1- and Ara h 2-specific IgE (**Table S2**). The analysis showed a substantial reduction in IgE-binding to PN proteins overall, and to Ara h 1 and 2 in particular (**Figure 4G–H**). To precisely define the impact of microbial metabolism on main PN allergens, we mono-sensitized mice to native Ara h 1 or recombinant Ara h 2 and used their sera to analyze IgE recognition of the digested PN fragments via Western blotting (**Figure 4I–J**). *Rothia* digestion of PN reduced IgE-binding to Ara h 1, most notably by strain R3. Similarly, *Staphylococcus* strains (S1 and S3) completely degraded and reduced IgE-binding to Ara h 1, though some residual fragments retained IgE-binding capacity. *Staphylococcus* strains with lower PN-degrading capacity, such as S2, showed less efficacy in reducing binding to Ara h 1 (**Figure 4I)**. On the other hand, sera from mice sensitized with recombinant Ara h 2 predominantly recognized a band around ⁓37 kDa, likely representing its oligomerized forms. Nevertheless, PN-digestion by *Rothia* strains impaired Ara h 2 recognition by IgE, particularly with strain R3. *Staphylococcus* strains S1 and S3 also reduced IgE-binding to Ara h 2, while strain S2 showed little effect (**Figure 4J)**. Altogether, these findings demonstrate the potential of *Rothia* and *Staphylococcus* strains to modify the structure of key PN allergens, including IgE-binding epitopes, thereby altering their recognition by both murine and human IgE.

**Figure 4.**
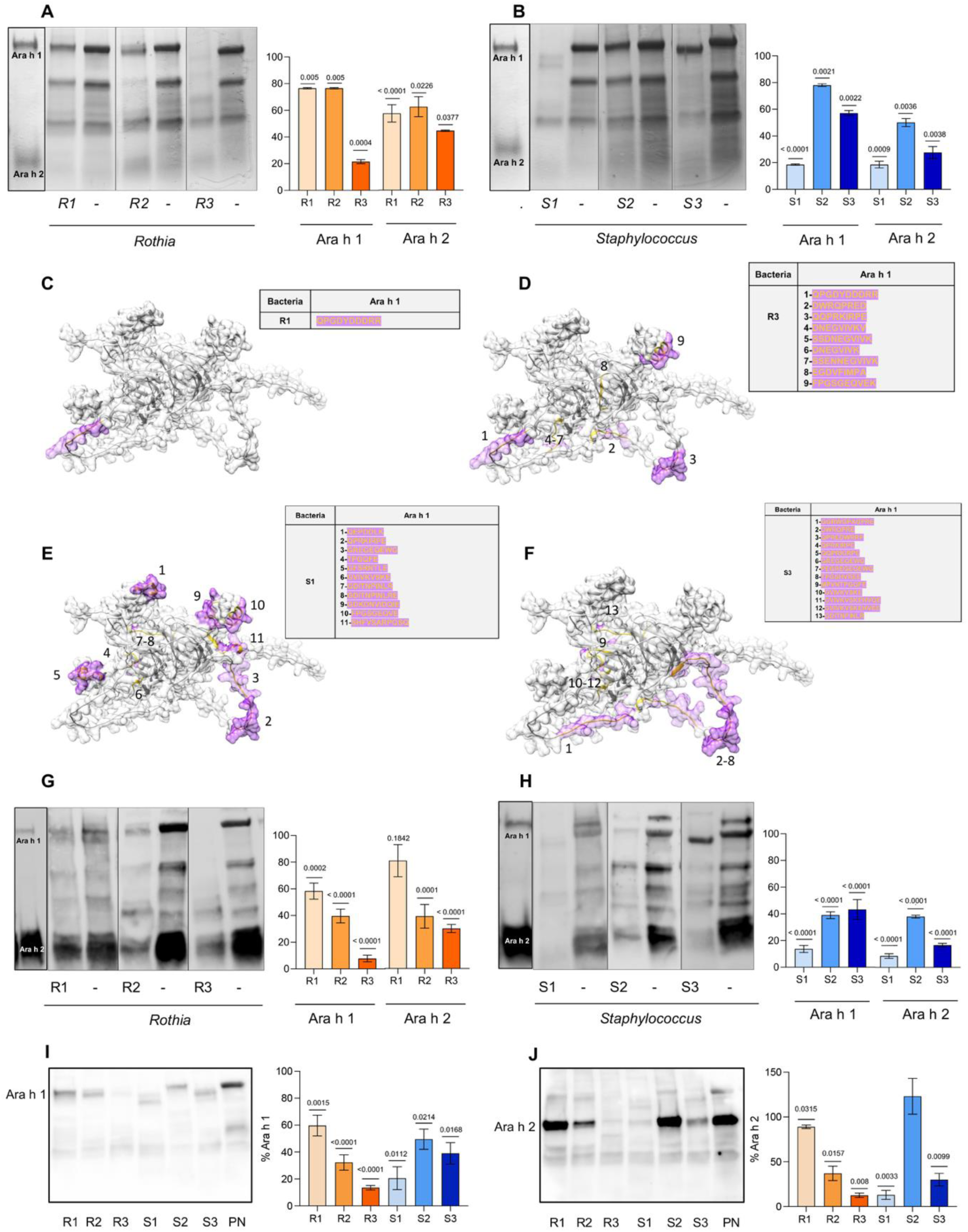
Peanut allergens are degraded by *Rothia* and *Staphylococcus*. Selected *Rothia* (R1–*R. aeria*, R2–*R. dentacariosa*, and R3–*R. mucilaginosa*) and *Staphylococcus* (S1–*S. epidermidis*, S2–*S. aureus*, and S3–*S. aureus*) strains were incubated with crude peanut extract (CPE) in liquid media to assess Ara h 1 and 2 degradation. (A–B) SDS-PAGE of Ara h 1 (64 kDa) and Ara h 2 (17 kDa) degradation by *Rothia* (A) and *Staphylococcus* (B) vs. controls. (C–F) Molecular modeling of Ara h 1 (Alphafold database code: AF-P43238-F1): peptides <3 kDa identified by mass spectrometry after digestion by R1 (C), R3 (D), S1 (E), and S3 (F) in yellow; human IgE-recognized epitopes in purple. (G–H) Western blot showing peanut (PN)-allergic human IgE-binding to PN allergens after *Rothia* (G) and *Staphylococcus* (H) digestion relative to controls. (I–J) Western blot showing IgE-binding from Ara h 1 (I) or Ara h 2-sensitized (J) mice after bacterial digestion. Bar charts show allergen quantification after bacterial digestion (mean ± SEM). *P* values calculated using unpaired *t*-test between undigested and bacterially-digested PN. n=4–6 (A, B, G–J).

### Microbial metabolism mitigates peanut allergenicity

The structural modifications of Ara h 1 and 2 caused by microbial metabolism affected IgE recognition, prompting us to assess the functional consequences using a mast cell activation assay^47,48^. We sensitized bone marrow-derived mast cells (BMMCs) with sera from mice allergic to either native Ara h 1 or recombinant Ara h 2 and assessed IgE-mediated mast cell activation after challenge with bacterial PN digests. This was done by challenging the BMMCs with either bacterially digested PN or undigested PN, and measuring CD63 and CD107a surface expression, along with β-hexosaminidase activity in the cell supernatants (**Figure 5 & Figure S6A**). Our initial controls confirmed that serum from Ara h 1-allergic mice induced BMMC activation following Ara h 1 or CPE challenge, but not upon Ara h 2 stimulation, and vice versa (**Figure S6B–C**). We then observed that, among the three *Rothia* strains, PN-digestion by strain R2 consistently reduced BMMC activation for both Ara h 1 and 2; while PN-digestion by strains R1 and R3 significantly decreased BMMC responses in Ara h 2-sensitized cells (**Figure 5A–B**). On the other hand, among the three *Staphylococcus* strains, PN-digestion by strain S1 significantly reduced BMMC activation in Ara h 1-sensitized cells **(Figure 5C)**, while BMMC responses to Ara h 2 were impaired by the digestion of the three strains (**Figure 5D**). In sum, these results demonstrate that the metabolism of key PN allergens by human oral bacteria can modulate the ability of these allergens to trigger IgE-mediated mast cell activation.

**Figure 5.**
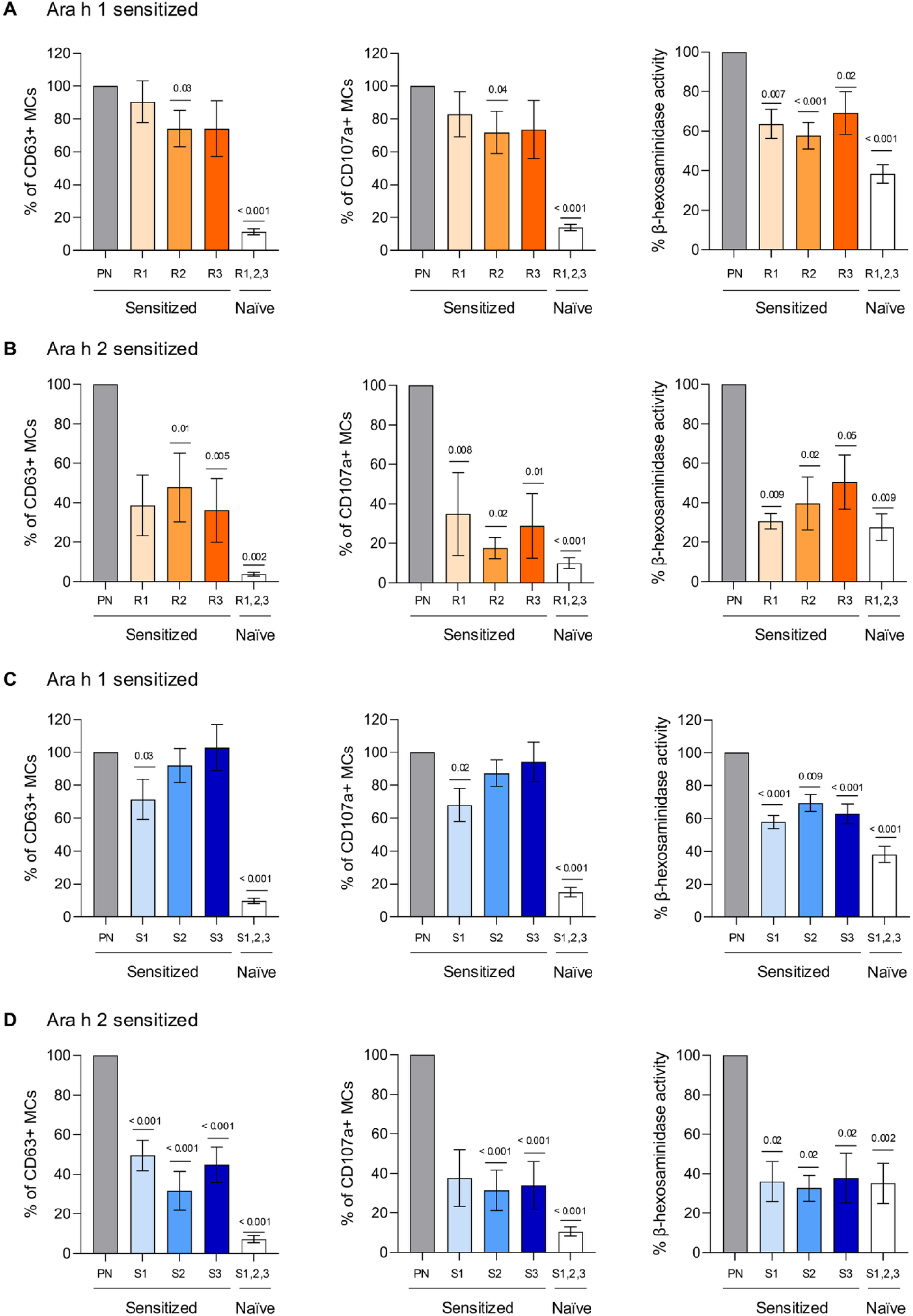
Challenge with bacterially digested peanut impairs mast cell activation. Mast cells (MCs) were sensitized with pooled sera from mice allergic to Ara h 1 or 2 and challenged with undigested peanut (PN; gray, positive control) or PN digested by *Rothia* (orange; R1*–R. aeria*, R2*–R. dentocariosa*, R3*–R. mucilaginosa*; n=3–5; A–B) or *Staphylococcus* (blue; S1–*S. epidermidis*, S2–*S. aureus*, S3–*S. aureus*; n=4–6; C–D). Negative controls (white) were MCs sensitized with sera from non-allergic mice and challenged with pooled bacterially-digested PN from all three *Rothia* or *Staphylococcus* species. Bar graphs indicate CD63 and CD107a expression, and β-hexosaminidase activity, normalized to 100%. Data = mean ± SEM. All three *Rothia* and *Staphylococcus* strains are displayed in the same graphs, but statistical analyses were performed separately for each strain vs. undigested PN. *P* values calculated using one-way ANOVA with Dunnett’s post-hoc test, except for specific cases R1-CD63-Ara h 2 (B), S2-CD107a-Ara h 1 (C), and S1-CD107a-Ara h 2 (D), where the Kruskal-Wallis test with Dunn’s correction was used. Comparisons with undigested PN used one-way ANOVA with Dunnett’s test for β-hexosaminidase activity (D), CD63 (A), and CD107a (A–C), or the Kruskal-Wallis with Dunn’s correction for β-hexosaminidase activity (A–C), CD63 (B–D), and CD107a (D). See also Figures S6–S8.

To further characterize the mechanisms underlying microbial allergen degradation, we performed whole-genome sequencing of the different *Rothia* and *Staphylococcus* species and identified candidate genes encoding protease repertoires (**Figure S7 & Figure S8**). Notably, conserved proteases in *Rothia* include members of the peptidase S8 family of subtilisin-like serine proteases that are secreted extracellularly, which have been previously attributed to degrading recalcitrant dietary proteins such as gluten^49-51^, and may likewise contribute to PN allergen degradation. Protease-encoding genes were more variable in *Staphylococcus*, with distinct, non-conserved extracellular proteases identified in each strain. When comparing PN-degrading functionality with annotated protease genes, we observed that potential extracellular proteases including the cysteine protease EcpA, were present in the most efficient degraders of immunodominant allergens. These results confirm the presence of proteolytic machinery potentially involved in PN degradation by bacteria.

### Microbial peanut metabolism alters systemic allergen access

To study the *in vivo* capacity of human oral bacteria to degrade PN allergens, we colonized GF C57BL/6 mice with bacterial strains exhibiting varying allergen-degrading capacities. We selected three *Staphylococcus* strains based on their *in vitro* PN-degrading capacities: one with strong capacity to degrade both Ara h 1 and 2 (S1), one that degrades only Ara h 1 (S3), and one with no detectable allergen-degrading capacity (S2). Additionally, we included a *Rothia* strain (R3) capable of degrading both Ara h 1 and 2. GF mice were used as controls. After colonization, mice were gavaged with PN (**Figure 6A**), as previously described for SPF and MM mice. Successful transfer of PN-degrading capacity was confirmed in colonized mice (**Figure 6B**). Notably, we observed a reduction in Ara h 1 and 2 levels in the small intestinal content of mice colonized with allergen-degrading bacteria, such as S1 and R3 (**Figure 6C**). Additionally, systemic levels of Ara h 1 and 2 were reduced in these mice (**Figure 6D**). In contrast, Ara h 1—and particularly Ara h 2—showed increased systemic levels in mice colonized with PN-degrading bacteria that had impaired allergen-degrading capacity (S2) (**Figure 6D**). To study whether bacterial metabolism alters allergen passage through the mucosa and subsequent systemic access, we assessed allergen translocation across *ex vivo* mouse small intestinal tissue using Ussing chambers. CPE, pre-incubated with or without the different bacterial strains was applied to the mucosal side of the chamber, and Ara h 1 and 2 levels were quantified on the serosal side of the tissue after 2 hours. We found that PN-degrading bacteria with impaired allergen-degrading capacity facilitated the passage of Ara h 1 and 2 through the intestinal mucosa (**Figure 6E**). These findings suggest that bacteria metabolize PN *in vivo* and control allergen passage through the intestinal mucosal barrier.

**Figure 6.**
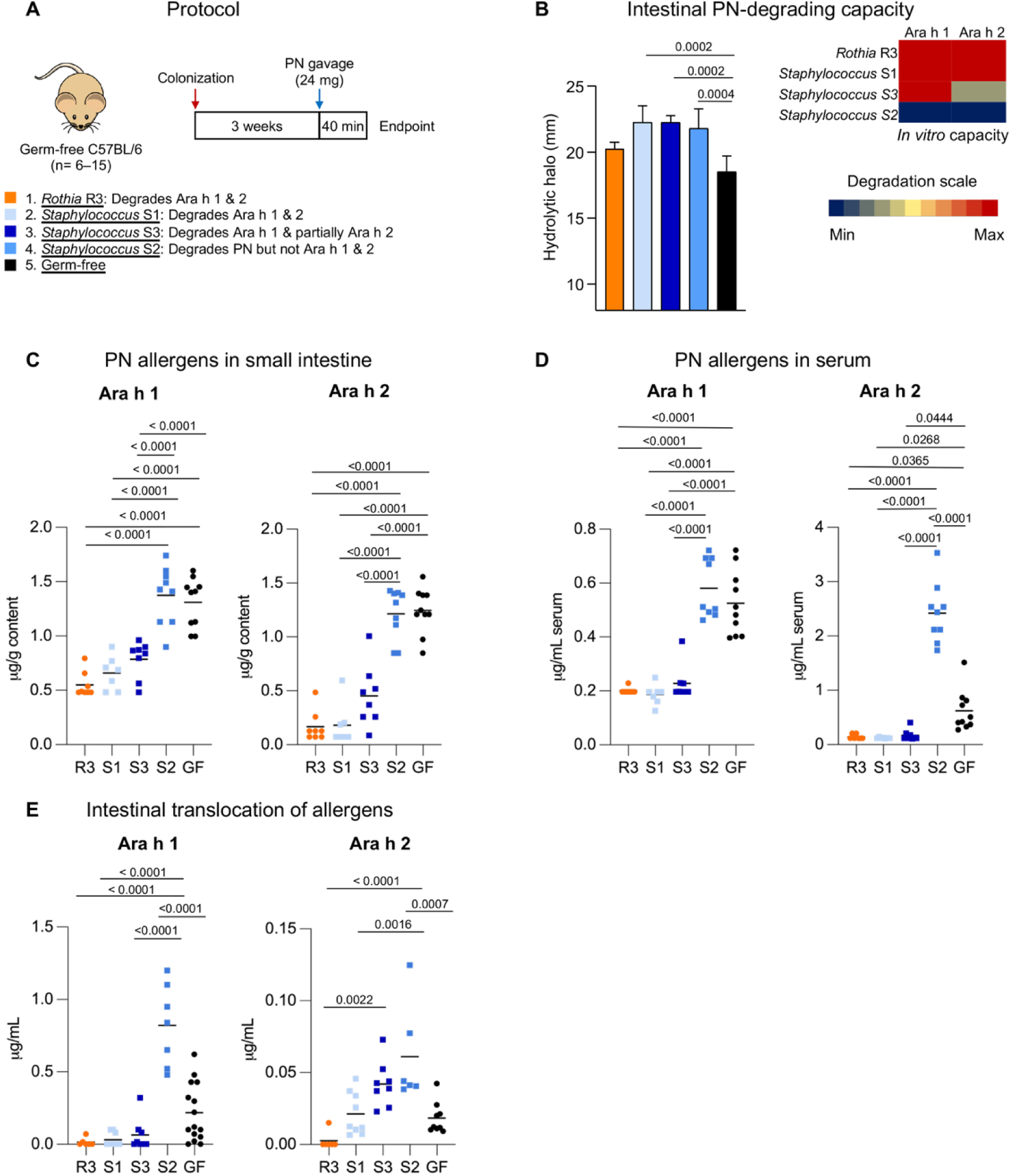
Bacteria dictate systemic access of allergens *in vivo*. (A) Experimental design. Germ-free (GF) C57BL/6 mice were mono-colonized with bacteria (R3–*Rothia mucilaginosa*, S1–*Staphylococcus epidermidis*, S3–*Staphylococcus aureus*, and S2–*Staphylococcus aureus*) for 3 weeks before peanut (PN) gavage; GF mice served as controls. Mice were sacrificed 40 min after PN gavage. (B) PN-degrading capacity of small intestinal contents measured by hydrolytic halo diameter on PN-agar, and degradation of Ara h 1 and 2 (heatmap, blue=0%, red=100% relative to crude peanut extract). (C–D) Ara h 1 and 2 in small intestinal content (C) and serum (D). (E) Translocation of PN allergens digested by bacteria from the mucosal to serosal side of small intestinal tissue in Ussing chambers (R3, orange; S1, light blue; S3, dark blue, S2, medium blue; non-growth control, GF, black). Data are shown as mean + SD (B) and mean where each dot = one mouse (C, D, E). n=7–10 mice/group (B–D) and n=6–15 mice/group (E); pooled from 2 independent experiments. *P* values calculated using a one-way ANOVA with Tukey’s post-hoc test.

### Microbial peanut metabolism dictates anaphylactic reactions *in vivo*

To evaluate the impact of microbial PN metabolism on anaphylactic responses, MM C3H/HeN mice—which have impaired microbial PN metabolism and are susceptible to anaphylactic reactions upon oral exposure—were sensitized to PN via i.p. injection and subsequently challenged with bacterially pre-digested PN (**Figure 7A**). Four groups of mice received an i.g. PN-challenge with either untreated PN (control) or PN pre-digested by one of the following bacteria: a) *Rothia* R3, b) *Staphylococcus* S1, or c) *Staphylococcus* S2, each exhibiting distinct allergen-degrading capacities. While all groups showed similar PN-specific IgE levels after sensitization (**Figure 7B**), serum mMCP-1 levels were lower in mice challenged with PN digested by *Rothia* R3 and *Staphylococcus* S1 (**Figure 7C**). Conversely, mice challenged with PN pre-digested by *Staphylococcus* S2, which has limited allergen-reducing capacity, exhibited higher serum levels of mMCP-1. To evaluate the protective capacity of *Rothia* colonization against food-induced anaphylaxis, MM C3H/HeN mice were sensitized to PN (i.g.) and then colonized with *Rothia* R3 with efficient PN-degrading capacity (**Figure 7D**). After sensitization, the mice were challenged to PN i.g. Despite having similar PN-specific IgE levels in serum and small intestinal mucosal mast cells (**Figure 7E & Figure S9A–E**), mice colonized with *Rothia* R3 demonstrated significantly reduced serum levels of mMCP-1 (**Figure 7F**), hypothermia (**Figure 7G**), and serum PAF-AH (**Figure S9B**). Furthermore, PN-challenge in MM mice colonized with or without *Rothia* after i.p. sensitization led to reduced mast cell degranulation and hypothermia in *Rothia*-colonized mice despite similar PN-specific IgE levels and mucosal mast cells (**Figure S9F–K**). Overall, these findings highlight the critical role of microbial allergen metabolism in modulating allergic responses *in vivo*.

**Figure 7.**
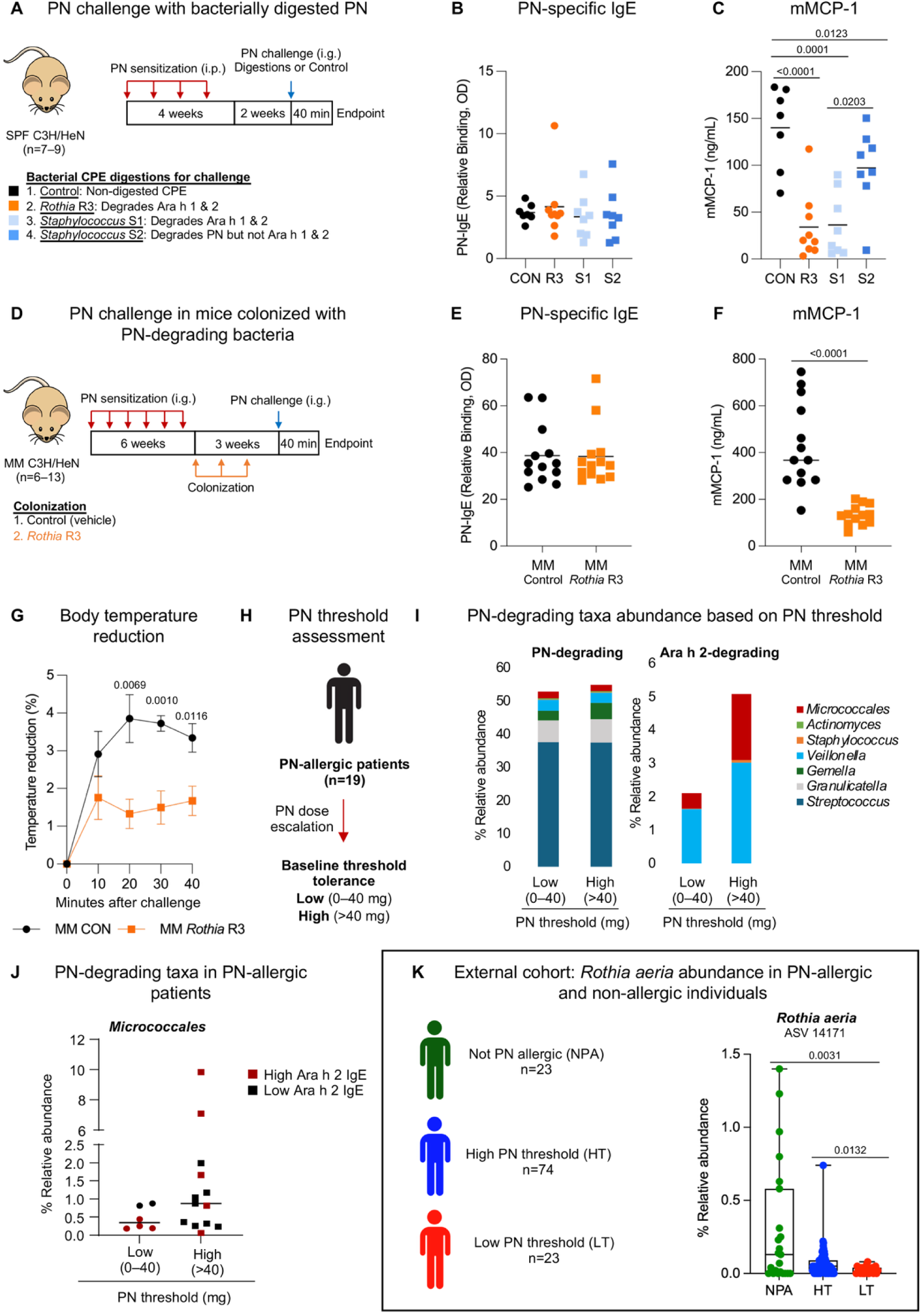
Microbial peanut metabolism reduces allergic reactions *in vivo* and is associated with allergen tolerance. (A) Experimental design. Specific pathogen-free (SPF) C3H/HeN mice were sensitized intraperitoneally (i.p.) to peanut (PN) weekly for 4 weeks and challenged intragastrically (i.g.) after 7 weeks with PN digested by *Rothia mucilaginosa* (R3), *Staphylococcus epidermidis* (S1), and *Staphylococcus aureus* (S2), or undigested PN. Mice were sacrificed 40 min later. n=7–9 mice/group; pooled from 2 independent experiments. (B) Serum PN-specific IgE before challenge. (C) Serum mucosal mast cell protease 1 (mMCP-1) after challenge. (D) Experimental design. Minimal microbiota (MM) C3H/HeN mice were sensitized i.g. to PN weekly for 6 weeks, then colonized with *Rothia* R3 weekly for 3 weeks. PN-challenge was given i.g. 3 weeks after the first colonization, followed by sacrifice 40 min later. (E) Serum PN-specific IgE before PN-challenge. n=13 mice/group; pooled from 3 independent experiments. (F) Serum mMCP-1 after PN-challenge. n=13 mice/group; pooled from 3 independent experiments. (G) Core body temperature reduction after PN-challenge. n=6 mice/group pooled from two independent experiments. (H) Experimental design. PN-allergic patients (n=19) were stratified by PN threshold (low –0-40 mg PN; high–>40 mg PN) prior to oral immunotherapy. (I) Relative abundance (%) of PN- and Ara h 2-degrading bacteria in low vs. high threshold patients. (J) Relative abundance of *Micrococalles* in the saliva of low vs. high PN threshold patients. Dot color represents high (≥20 kUA/mL, red) and low (<20 kUA/mL, black) serum Ara h 2-IgE. (K) *Rothia aeria* ASV 14171 abundance in saliva from 16S rRNA data of an external cohort of 120 children (23 non-PN-allergic controls (NPA), 74 PN-allergic with high PN threshold (HT; >443 mg), and 23 PN-allergic with low PN threshold (LT; <443 mg)) who underwent double-blind, placebo-controlled PN-challenges. Data are presented as mean with dots for individual mice or humans (B, C, E, F, J), as mean ± SEM (G), or as interquartile range with whiskers extending to min/max (K). *P* values: one-way ANOVA with Tukey’s post-hoc test (C), unpaired Student’s *t*-test (E–G), and Kruskal-Wallis test with Dunn’s post-hoc test (K). See also Figures S9 and S10.

### Peanut-degrading bacteria are increased in allergic patients better tolerating peanut

To investigate the role of the oral microbiota in allergic reactions, we collected saliva samples from a small well-defined cohort of PN-allergic patients (**Figure 7H & Table S1)**. Prior to initiating oral immunotherapy (OIT), allergic patients underwent a PN-challenge to determine their threshold for reactivity (PN threshold). We analyzed serum levels of IgE and the composition of the oral microbiota. Notably, serum levels of IgE specific to Ara h 1 and 2 showed a poor association with PN threshold **(Figure S10A)**, suggesting the role of additional factors. Examining the oral microbiota, we observed non-significant clustering based on PN threshold (**Figure S10B)**. While PN-degrading taxa were present across patients with varying PN thresholds, individuals with higher PN thresholds exhibited a greater abundance of taxa capable of degrading Ara h 2 **(Figure 7I)**. Notably, *Micrococcales*—a taxon that includes *Rothia* and *Micrococcus*, both efficient at degrading immunodominant Ara h 1 and 2—was reduced in PN-allergic patients with low PN threshold (0–40 mg), independent of serum Ara h 2-specific IgE levels (**Figure 7J**).

Interestingly, some high-threshold patients with elevated Ara h 2-specific IgE had increased *Micrococcales*, potentially implicating non-IgE factors in reactivity. To validate our clinical findings, we examined the relative abundance of *Rothia* in saliva of PN-allergic and non-allergic children in an external cohort of 120 children who underwent double-blind, placebo-controlled food challenges^52^. The participants included 23 children with no allergy (controls), 74 with high-threshold PN allergy (reacting to ≥ 443 mg cumulative PN protein), and 23 with low-threshold PN allergy (reacting to <443 mg cumulative PN protein). Notably, *Rothia aeria* ASV14171 was significantly more abundant in the saliva of non-PN-allergic and high PN threshold allergic individuals compared to allergic individuals with low PN thresholds (**Figure 7K**). Other *Rothia* species also showed a trend toward enrichment in the groups (**Figure S10C**). These findings suggest that the oral microbiota could serve as a predictive marker of threshold reactivity to PN, highlighting the potential importance of microbial allergen metabolism in IgE-mediated reactions.

## DISCUSSION

The human microbiome is essential for maintaining homeostasis, and disruptions in its composition are associated with immune disorders. The oro-gastrointestinal microbiota has gained attention for its role in influencing oral tolerance and sensitization to food antigens^18-20,22,53^. While studies have reported differences in the intestinal microbiota composition of food-allergic patients^23-25^, the microbial mechanisms underlying food-induced anaphylaxis remain largely unknown. In this study, we show that human saliva and small intestinal content harbor allergen-degrading bacteria capable of metabolizing immunodominant PN allergens, thereby modulating IgE-mediated reactions to foods. Identifying the microbes involved in PN metabolism in humans and characterizing the related microbially-mediated IgE-specific immune responses could have implications for reducing the severity of allergic reactions.

The antigenic properties of many foods stem from their resistance to gastrointestinal digestion^28,29^. PN allergens, for example, are poorly metabolized by human digestive enzymes^26,27^, whereas the intestinal microbiota encodes diverse metabolic pathways absent in the host that participate in food digestion^54^. For example, the capacity of the intestinal microbiota to degrade other recalcitrant antigenic proteins, such as gluten^55^, has been demonstrated^56,57^. Using gnotobiotic mouse models, we demonstrate that microbiota composition and complexity contribute to PN allergen metabolism *in vivo* and the amount of allergen reaching the intestine and circulation. While the intestinal microbiota affects host metabolism and food-dependent immune responses through mechanisms such as altering the intestinal barrier, motility, or immune activation^18,58-60^, we confirmed the presence of PN-degrading bacteria capable of digesting Ara h 1 and 2 along the gastrointestinal tract in wild-type SPF mice. These findings reveal the role of the intestinal microbiota in allergen metabolism and raise important questions about how diet-microbiota interactions shape food allergy.

Food allergies are primarily IgE-mediated^61^, and even minute allergen doses can trigger life-threatening reactions ^62^. We hypothesized that microbial metabolism modulates IgE-dependent responses to PN. First, we demonstrated that mice with limited PN-degrading capacity (MM) developed enhanced allergic reactions to PN than those with efficient PN metabolism (SPF) following i.g. sensitization and challenge. MM mice exhibited greater mucosal mast cell activation, likely increasing intestinal permeability and facilitating systemic allergen access^63,64^—a critical step in IgE-mediated anaphylaxis^65^. Intriguingly, MM mice also developed higher titers of antigen-specific IgE than SPF controls, potentially due to their reduced ability to metabolize PN allergens. Similar increases in antigen-specific IgE have been reported in GF mice and mice colonized with limited or allergic-associated microbial communities following sensitization^31,66-68^. Given that food-induced anaphylaxis is largely IgE-dependent^69,70^, and the microbiota may influence IgE-mediated food sensitization independently of allergen degradation^18^, differences in sensitization between MM and SPF microbiota could explain the severity of allergic reactions. To bypass local microbial factors, we evaluated the role of microbial metabolism in IgE-mediated food responses using additional models of systemic^71^ and passive^48,72^ sensitization. Following PN-challenge, MM mice with limited PN-degrading capacity developed enhanced allergic reactions, characterized by increased mast cell degranulation and hypothermia compared to conventional SPF mice, despite similar PN-IgE titers. Altogether, these results indicate that microbial metabolism is a key determinant of IgE-mediated food allergy in mice.

Food-induced anaphylaxis usually results from accidental allergen exposure, posing a persistent threat that significantly impacts quality of life^73,74^. Although co-factors such as exercise, estrogen levels, alcohol, and drugs can heighten anaphylaxis risk^14,75^, variations in allergen thresholds and susceptibility remain poorly understood. A recent study demonstrated that anaphylaxis severity is independent of the amount of allergen ingested, emphasizing individual sensitivity and the potential role of microbiota in modulating IgE-responses^76^. To explore the potential role of microbial allergen metabolism in humans, we searched for PN-degrading bacteria in healthy donors, focusing on the oral cavity and small intestine where ingested food first interacts with the microbiota. While most microbiota research has centered on the large intestinal lumen due to its ease of sampling and high bacterial density, recent studies have highlighted the stability and complexity of the oral and small intestinal microbiota, as well as their relevance in health and disease^77^. Moreover, the oral microbiota serves as a continuous source of bacteria for the intestinal tract, seeding the small intestine with taxa such as *Rothia* and *Staphylococcus^78^*, which may persist and contribute to allergen metabolism as food moves through the upper gut^79^. Using SDS-PAGE, Western blotting, and proteomics, we confirmed the presence of PN-degrading bacteria in the oral cavity with strain-specific capacities to degrade immunodominant PN allergens, disrupt human IgE epitopes, and alter IgE-binding of sera from PN-allergic patients and PN-sensitized mice. Overall, these results confirm the capacity of bacteria to potentially alter allergenicity in the oral cavity and small intestine of humans.

Dominant taxa in the oral cavity and small intestine such as *Rothia*, and related genera like *Micrococcus*, consistently digested both Ara h 1 and 2 *in vitro,* altering IgE recognition. The capacity of *Rothia* to degrade other dietary antigens such as gluten proteins has been also suggested^51,80^. Remarkably, impaired IgE recognition of PN upon *Rothia* digestion hampered the activation of the classical pathway of anaphylaxis both *in vitro* (mast cell assays) and *in vivo* (PN-allergic mice challenged with digested PN). In addition, the PN-degrading capacity of *Rothia* was validated *in vivo*, reducing the concentration of major PN allergens reaching the small intestine and circulation. We also tested the capacity of *Rothia* strains to alter IgE-mediated reactions following colonization in a competitive environment. To avoid changes in the adaptive immune response, MM mice were colonized following either i.g. or i.p. sensitization. MM mice colonized with *Rothia* had reduced IgE-mediated allergic responses and mucosal mast cell degranulation following challenge compared to MM controls. These results confirm that the protective effects of allergen metabolism arise by reducing allergen availability in the intestinal environment rather than by altering systemic sensitization. Altogether, our data demonstrate that PN-degrading bacteria can modulate acute allergic responses *in vivo*.

Microbial PN metabolism does not always protect against IgE-mediated responses. Certain PN-degrading genera, such as *Gemella* or *Streptococcus,* lacked allergen-degrading capacity *in vitro*, while *Staphylococcus*, a main PN-degrading taxon in the oral cavity, displayed strain-specific effects in our work. Others have reported that skin colonization by *Staphylococcus aureus* induces keratinocyte release of IL-36α^81^, a pro-inflammatory cytokine linked to allergic sensitization to foods^33^. Additionally, *Staphylococcus* and other microbial proteases such as EcpA can induce immune activation and low-grade inflammation in the colon^82^, and promote inflammatory responses to innocuous antigens^83,84^. Our findings revealed that *Staphylococcus* strains vary in their capacity to remove PN allergens and alter IgE-mediated recognition and mast cell activation *in vitro*. In gnotobiotic mice colonized with *Staphylococcus* strains exhibiting differing PN-degrading capacities, efficient strains reduced Ara h 1 and 2 levels in the small intestine and serum. However, these immunodominant peptides increased systemically in mice colonized with a strain lacking efficient PN-degrading capacity. Enhanced passage of Ara h 1 and 2 through the intestinal mucosa after partial digestion of PN by this *Staphylococcus* strain was also confirmed with Ussing chambers, highlighting the role of bacterial activity in systemic allergen access, a critical step in anaphylactic reactions^65^. These results indicate that *Staphylococcus* species, including *Staphylococcus aureus,* may facilitate allergen passage through the mucosal barrier and promote allergic inflammation when allergen degradation is incomplete^85,86^. Some *Staphylococcus* strains can also compromise the epithelial barrier and increase permeability directly^87,88^. Therefore, microbes may have a dual impact on IgE-mediated immune responses, depending on the bacteria’s ability to degrade PN allergens, regardless of its taxonomy. Complete microbial degradation of PN allergens reduces IgE responses, while inefficient metabolism may promote anaphylaxis by enhancing the passage of allergens through the mucosa. Altogether, the ability of the microbiota to completely degrade immunodominant allergens and reduce IgE recognition of PN significantly influences subsequent mast cell activation and anaphylactic reactions.

Despite these findings, the precise microbial proteases responsible for PN allergen metabolism remain to be fully identified. Whole-genome sequencing of different *Rothia* species revealed diverse protease repertoires, including subtilisin-like serine proteases. Members of this family have been shown to degrade other recalcitrant dietary proteins^49-51^, such as gluten, suggesting they may contribute to PN allergen metabolism. Efficient PN-degrading *Staphylococcus* possessed annotated extracellular proteases, including the virulence-associated cysteine protease EcpA^89,90^. However, given the broad proteolytic capacity of the oral microbiota, multiple enzymes could be involved in modulating allergenicity. Future studies integrating genetic modulation will be required to pinpoint the specific microbial enzymes mediating these effects.

While our findings consistently support the role of microbial metabolism in modulating PN allergenicity, variability was observed in our results across models and microbial communities. Both technical (*e.g.*, antibody and *ex vivo* assay variability) and biological factors (*e.g.*, postprandial state, vendor-specific SPF microbiota) likely contributed. We used i.g. challenges to focus on mucosal mast cell activation, though these may induce weaker clinical anaphylaxis than the i.p. route and add variability. Specifically, immune responses upon i.g. challenge are influenced by factors including allergen delivery, intestinal permeability, gastric emptying, and genetic background. This variability is further amplified after i.g. sensitization that also enhances mast cell recruitment to the small intestinal mucosa and intestinal permeability^91^. GF mice, the gold standard for studying microbially-mediated phenotypes, were used to investigate PN metabolism, but they exhibit altered intestinal barrier function, impaired motility, and increased proteolytic activity, and their underdeveloped immunity can exaggerate sensitization and anaphylaxis^66,68,92-94^. Importantly, sensitization was not the primary focus of this study and allergic responses may be influenced by factors beyond microbial^22^, such as the extent of mast cell recruitment to the intestine^91^. Furthermore, leukotriene synthesis which has been shown to differ by mouse strain, also affect allergen uptake in the gastrointestinal tract and subsequently oral anaphylaxis^3,4^. To better isolate microbial effects, we used MM mice, which provide minimal but stable microbiota with limited endogenous PN-degrading activity. MM mice serve as a stronger comparator for high PN-degrading capacity microbiota or as a model for supplementation with PN-degrading taxa. Colonization of MM mice with PN-degrading *Rothia* after i.g. or i.p. sensitization to PN reduced anaphylaxis markers and mast cell degranulation. Microbial composition and complexity were key determinants of allergen degradation and IgE-mediated responses, as shown by a negative correlation between alpha-diversity and both intestinal allergen levels and serum mMCP-1 in MM and SPF mice. This suggests that variability in PN degradation and anaphylaxis observed in our study may be partially attributed to individual differences in microbiota composition and complexity, with some SPF mice potentially harboring fewer PN-degrading bacteria. Importantly, the PN-degrading taxa identified in humans, such as *Rothia*, are largely absent in SPF mice, and as a result these correlations between diversity and intestinal allergen levels reflect broader ecological relationships between microbiota complexity and PN-degradation, rather than the contribution of these specific human-derived taxa. Accordingly, the functional relevance of *Rothia* and *Staphylococcus* was evaluated separately using defined colonization models in this study, allowing mechanistic testing of these human PN-degrading isolates. Together, these complementary approaches—ecological associations in murine communities and taxa-specific experiments in gnotobiotic models—provide coherent and convergent evidence for microbial control of PN metabolism and anaphylaxis.

The primary recommendation for food allergic patients is strict avoidance of allergens, yet this is a challenge for ubiquitous foods such as egg, wheat, milk or PN, leading to frequent accidental exposures^10,12,95,96^. Allergen thresholds differ among PN-allergic patients, with some individuals experiencing anaphylaxis to trace amounts^97,98^. To investigate the role of microbial allergen metabolism in food-induced anaphylaxis, we characterized the oral microbiota of PN-allergic patients initiating OIT with varying allergen thresholds. Allergen-specific IgE did not correlate well with allergen thresholds in our study, consistent with previous findings^99,100^. However, patients with higher PN thresholds showed greater abundance of *Micrococcales* (an order including PN-degrading *Rothia* and *Micrococcus*) than highly sensitive PN-allergic patients, independent of IgE titers. Consistent with this observation, our analysis of an external pediatric cohort revealed that *Rothia aeria* was significantly more abundant in the saliva of non-PN-allergic and high-threshold PN-allergic individuals compared to PN-allergic individuals with low thresholds, with other *Rothia* species also showing a trend toward enrichment^52^. A previous study also reported an increased relative abundance of *Rothia* in the saliva of PN-allergic children after undergoing OIT^101^, and *Veillonella*, an oral taxon with PN-degrading capacity, is increased in those with greater reaction thresholds^102^. Together, these findings suggest that allergen threshold associates with the metabolic activity of the oral microbiota against food allergens, and may serve as a marker of reactivity. Still, the mechanistic link between oral microbiota and allergen threshold in humans remains speculative, requiring preclinical and longitudinal validation. Longitudinal assessment of oral microbiota and allergen reactivity during PN OIT in future studies will further inform how microbiota may influence OIT outcomes. Although OIT remains the only allergen-specific treatment with disease-modifying potential, it is associated with limitations such as events that compromise patient adherence, quality of life, and overall effectiveness^74,103,104^. Functional analysis of the oral microbiota may help identify patients with low allergen thresholds less suitable for OIT, while harnessing PN-degrading bacteria may raise allergen thresholds. Characterizing PN-degrading bacteria could also be applied to reduce cross-contamination risks—a leading cause of unintentional allergen exposure and a significant concern in food allergy management^96,105-107^—and improve the formulation of hypoallergenic products. In summary, our findings reveal a novel microbial mechanism of relevance in food-induced anaphylaxis. Our results underscore the role of the human microbiota in dictating the severity of IgE-mediated reactions through allergen metabolism and highlight the therapeutic potential of harnessing bacterial allergen-degrading capabilities for managing food allergies.

## RESOURCE AVAILABILITY

### Lead contact

Further information and requests for resources and reagents should be directed to the lead contact Alberto Caminero.

### Materials availability

This study did not generate new unique reagents.

### Data and code availability

The microbiome sequencing data generated in this study have been deposited in the NCBI BioProject database under the accession number PRJNA1223483. The mass spectrometry proteomics data have been deposited to the ProteomeXchange Consortium via the PRIDE partner repository with the dataset identifier PXD060888 and 10.6019/PXD060888. Any additional information required to reanalyze the data reported in this paper is available from the lead contact upon reasonable request.

## ACKNOWLEDGMENTS

We thank the McMaster University AGU staff, Dr. Heather G. Galipeau, Joe Noratangelo, Michael Rosati, and Sarah Armstrong for their assistance with mouse care. We thank the McMaster Genome Facility, Laura Rosati, and Michelle Shah for technical support with 16S rRNA gene sequencing. We thank Gaston Rueda for assistance with clinical samples. We thank the staff of the Proteomics Unit at CNB-CSIC, particularly Dr. Alberto Paradela, for analysis and technical support. We are grateful to Drs. Pablo Rodríguez del Río, Salvador Iborra and Francisco Sánchez-Madrid for their critical review of the manuscript. This work was supported by grants from The Nutricia Research Foundation (NRF-2021-13) and the New Frontiers in Research Fund (NFRFE-2019-00083) to R.J.-S. and A.C.; and by a grant from the European Food Safety Authority (GP/EFSA/ENCO/2020/02-1) to F.J.M. R.J.-S.’s laboratory is supported by the Instituto de Salud Carlos III (ISCIII; CP20/00043; PI22/00236; PI25/00477; RD24/0007/0037), by the Ministerio de Ciencia, Innovación y Universidades (CNS2024-154194) and by the Community of Madrid (IND2024/BMD-33910). A.C. holds a Family Douglas Research Chair in Intestinal Research and is funded by the Canadian Institutes of Health Research (CIHR; Project Grant 202010PJT), NSERC (DGECR-2022-00221) and by Crohn’s and Colitis Canada (CCC, Grants-In-Aid grant). L.E.R. is supported by a Frederick Banting and Charles Best Canada Graduate Scholarship-Doctoral award from CIHR and a graduate scholarship from the Farncombe Institute. E.S.-M. is supported by a Formación de Profesorado Universitario grant (FPU23/03341) from Ministerio de Universidades. S.U.P. is supported by NIAID NIH R01 grants 1R01AI155630 and 1R01AI182001. L.M.-S. is supported by the Investigo program through the Ministerio de Trabajo y Economía Social, Servicio Público de Empleo (SEPE), which is funded by “Plan de Recuperación, Transformación y Resiliencia” and “NextGenerationEU” of the European Union (2022-C23.I01). C.L.S and E.N.-B are supported by a Río Hortega (CM23/0019) and a Sara Borrell grant (CD23/00125), respectively, from ISCIII. E.F.V. is supported by a CIHR Project Grant (PJT-183881) and holds a Tier 1 Canada Research Chair in Microbial Therapeutics and Nutrition in Gastroenterology. L.Z. and S.B. received funding from NIAID NIH R01 AI147028 and U196053.

## AUTHOR CONTRIBUTIONS

Conceptualization, R.J.-S., A.C.; Methodology, R.J.-S., A.C.; Investigation, E.S.-M., L.E.R., M.G.-R., B.B., D.H., P.H., D.C., G.Y., R.D., J.F.E.K., T.D.W., L.M.-S., C.L.S., E.N.-B., X.Y.W., G.D.P., L.Z., F.J.M., R.J.-S., A.C.; Resources, M.G.-A., A.D.-P., E.F.V., Y.R.C., P.O., F.V., C.B., J.F.E.K, M.J., W.G.S., S.U.P., F.J.M., M.G.S., S.B., P.B.; Writing – Original Draft Preparation, E.S.-M., L.E.R., R.J.-S., A.C.; Writing – Review & Editing Preparation, E.S.-M., L.E.R., R.J.-S., A.C.; Visualization Preparation, E.S.-M., L.E.R.; Supervision, R.J.-S., A.C.; funding Acquisition, R.J.-S., A.C.

## SUPPLEMENTAL INFORMATION INDEX

Figures S1-S10 and their legends, as well as Tables S1 and S2 can be found in Document S1.

## STAR PROTOCOLS

### RESOURCE AVAILABILITY

#### Lead contact

Requests for further information or resources should be directed to the lead contact Alberto Caminero.

#### Materials availability

This study did not generate new unique reagents.

#### Data and code availability

The microbiome sequencing data generated in this study have been deposited in the NCBI BioProject database under the accession number PRJNA1223483. The mass spectrometry proteomics data have been deposited to the ProteomeXchange Consortium via the PRIDE partner repository with the dataset identifier PXD060888 and 10.6019/PXD060888.

## EXPERIMENTAL MODEL AND STUDY PARTICIPANT DETAILS

### Human samples

Human saliva from peanut (PN) allergic patients entering oral immunotherapy (OIT; n=19) were collected. These samples were obtained from children aged 1-14 years who participated in a standard clinical oral food challenge to establish PN threshold prior to clinical management with PN OIT at Massachusetts General Hospital. Saliva was collected with the SalivaBio Children’s Swab (Salimetrics) following the manufacturer’s protocol, and samples were stored at -80°C until further use. PN- and antigen-specific Ig levels in plasma from patients undergoing OIT were measured using a Phadia ImmunoCAP 1000 instrument (ThermoFisher) according to the manufacturer’s instructions. Saliva samples from healthy volunteers (n=13) were collected at McMaster University under BUP#423. All patients and healthy volunteers provided written informed consent as per local institutional review board guidelines. Supportive and demographics data can be found in Table S1.

To complement oral sampling, small intestinal aspirate samples (mid-jejunal washes) were obtained from non-allergic control individuals without any chronic gastrointestinal or metabolic conditions (n=5). These samples were collected under an ongoing clinical protocol led by Dr. Premysl Bercik at McMaster University (HIREB #15311). Push enteroscopy was performed following an overnight fast, and jejunal washes (15–20 mL) were collected as previously described^78^. Briefly, sterile saline was applied using a water jet to dislodge mucus-associated and luminal bacteria from the mid-jejunum and samples were frozen with or without 20% glycerol for further use. Small intestinal microbiota was analyzed by 16S rRNA sequencing, and samples were cultured for isolation and identification of PN-degrading bacteria. Supportive and demographics data can be found in Table S1.

Human sera for Western blotting were provided by the Allergology Service at Hospital Universitario de La Princesa (Madrid, Spain). The Research Ethics Committee of Hospital Universitario de La Princesa approved the study protocol (reference 4460). All the donors provided written informed consent with no conflict of interest. Donor sera characterization is detailed in Table S2.

### Mice

Age-, sex-, and strain-matched controls were used in all experiments. Specific pathogen-free (SPF) C57BL/6 mice were purchased from Charles River and Taconic, while SPF C3H/HeN mice were purchased from Taconic. Germ-free (GF) C57BL/6 mice used for colonization experiments were originally purchased from Taconic and rendered GF by two-stage embryo transfer. GF mice were bred in flexible-film gnotobiotic isolators at McMaster University’s Farncombe Family Digestive Health Research Institute Axenic Gnotobiotic Unit (AGU).

C3H/HeN mice harboring minimal microbiota (MM) were bred and housed under gnotobiotic conditions at the AGU. These MM mice were originally derived from Altered Schaedler Flora (ASF) C3H/HeN mice purchased from Taconic and have been maintained at the AGU for over 20 generations. While the classical ASF consortium consists of eight defined bacterial strains^82,108^, long-term independent maintenance has resulted in minor divergence from the original composition. We therefore refer to these mice as harboring a MM. Importantly, this community remains highly restricted in taxonomic diversity and free of proteobacteria. Compositional data are presented in the manuscript.

All mice were maintained on a 12-h light-dark cycle and fed an autoclaved chow diet and sterile water *ad libitum*. Eight-to-twelve-week-old mice were used throughout. All procedures were approved by the Environmental Council of the Community of Madrid with PROEX references 45.2/20 and 9.7/23, and the McMaster University Animal Care Committee and McMaster University Animal Research Ethics Board in accordance with the Animal Utilization Protocol #22-26. Ethical regulations for animal research were strictly followed.

### Food allergy and anaphylaxis model

To study the impact of microbes on PN metabolism and anaphylaxis, C3H/HeN mice with different microbiota were sensitized to PN using distinct techniques. 1. Intra-gastric sensitization: mice were provided 3.25 mg of crude PN extract (CPE; Greer Laboratories Inc) and 0.625 µg of cholera toxin (List Biological Laboratories) in 0.1 mL of PBS by oral gavage once a week for 6 weeks; 2. Intra-peritoneal sensitization: mice were provided 3.25 mg of CPE and 6.25 µg of cholera toxin in 0.1 mL of PBS intra-peritoneally once a week for 4 weeks; 3. Passive sensitization-mice were subjected to 3 intra-peritoneal injections (0.1, 0.2, and 0.4 mL) of serum obtained from intra-peritoneally sensitized PN-allergic mice as described^48^. After sensitization, mice were challenged with a 24 mg bolus of CPE in 0.8 mL of PBS by oral gavage. In some experiments, CPE was pre-digested with PN allergen-degrading bacteria (*Rothia* R3, *Staphylococcus* S1, or S2), or mice were colonized with PN-degrading bacteria (*Rothia* R3), prior to the oral challenge to evaluate the impact of bacterial PN metabolism on allergenicity and anaphylaxis. Serum was collected 48 h before challenge. Mice were monitored for 40 min after challenge for changes in rectal (core) body temperature. Peripheral blood was collected, and mice were euthanized 40 min after the oral challenge. Mucosal mast cell protease 1 (mMCP-1) was quantified in serum after the PN-challenge, as it is a key biomarker of anaphylaxis in mice, serving a similar role to serum tryptase for indicating mast cell activation in humans^109^. Platelet-activating factor acetylhydrolase (PAF-AH) was quantified in serum as a marker of anaphylaxis.

### Microbiota model for studying microbial PN metabolism

To investigate the role of microbiota in PN metabolism, we used two colonization approaches: 1. We studied PN metabolism in mice colonized with MM and SPF microbiota, and GF controls; 2. GF C57BL/6 mice were mono-colonized with selected PN-degrading bacteria (*Rothia* R3 or *Staphylococcus* S1, S2, or S3). After colonization, mice were administered 24 mg of CPE in 0.8 mL of PBS by oral gavage. Small and large intestinal contents and serum from peripheral blood were collected 40 min post-administration for quantification of PN allergens Ara h 1 and 2 in biological samples by ELISA (Indoor Biotechnologies) as described below. Additionally, small intestinal contents were incubated *ex vivo* with CPE overnight (o/n), and remaining Ara h 1 and 2 levels were quantified by ELISA (Indoor Biotechnologies). C57BL/6 mice were used in these experiments because of difficulties generating C3H/HeN mice under GF conditions.

### Characterization of bacterial PN-degradation and immune activation

To assess bacterial degradation of PN allergens and its impact on immune activation, we performed *in vitro* digestions, proteomic characterization, and mast cell activation assays. Bacterial isolates were incubated with CPE and degradation of Ara h 1 and Ara h 2 was quantified by ELISA. Then, *Rothia* (R1, R2, and R3) and *Staphylococcus* (S1, S2, and S3) candidates were selected for further characterization by SDS-PAGE, and Western blotting. Western blotting was performed using sera from PN-allergic patients as well as sera from mice allergic to Ara h 1 or to Ara h 2, to identify IgE-binding epitopes in the digested PN allergens. Proteomic analysis using nano liquid chromatography coupled to mass spectroscopy was used to characterize the degradation products and PN-specific epitopes. For immune activation studies, bone marrow-derived mast cells (BMMCs) were cultured and sensitized to PN with sera from PN-allergic mice. These cells were challenged with bacterially digested PN to evaluate BMMC activation by β-hexosaminidase release and flow cytometry analysis of degranulation markers (CD63, CD107a)^47^.

## METHOD DETAILS

### Mouse colonization procedures

For colonization of GF C57BL/6 mice with MM and SPF microbiota, fresh cecal and colon contents were harvested from SPF C57BL/6 mice (Taconic) or MM C3H/HeN mice, diluted 1:10 in sterile PBS in anaerobic conditions, and 0.2 mL of each cecal/fecal suspension was orally gavaged to mice.

For mono-colonization of GF C57BL/6 mice with PN-degrading bacteria, *Rothia* R3 or *Staphylococcus* S1, S2, or S3 were grown o/n in brain heart infusion broth (BHI; Research products International Corp.) and 10^9^ colony forming units (CFU) from the culture were provided by oral gavage with PBS as a vehicle. GF controls were provided PBS by oral gavage.

For the PN anaphylaxis model, MM C3H/HeNTac mice received 10^9^ CFU total of *Rothia* R3, by oral gavage, administered once per week for three weeks after sensitization and before PN-challenge. Sterile PBS was used as a vehicle and served as a non-colonization control.

Colonization was confirmed by culturing intestinal content combined with PCR of the 16S rRNA gene and Sanger sequencing to verify bacterial identity (methods detailed below). These checks were performed throughout the experiments and at endpoint. Mice with evidence of contamination or failed colonization were excluded from the analysis. For selected experiments, 16S rRNA microbiota analysis was also performed to confirm overall microbial community composition.

### PN-specific IgE determination

Peripheral blood was collected by retro-orbital bleeding 48 h before the PN-challenge. Serum PN-specific IgE was measured using a sandwich ELISA adapted from previously described methods^48^. Briefly, 96-well plates (high-binding, flat-bottom, polystyrene; Corning, USA) were coated o/n at 4°C with 2 µg/mL rat anti-mouse IgE antibody diluted in PBS. Plates were washed with PBS/0.05% Tween-20 (PBS-T; Sigma Aldrich), then blocked with 1% BSA in PBS-T for 2 h at room temperature (RT). Plates were washed thoroughly with PBS-T, then serum samples were incubated on the plate at 1/2 dilution o/n at 4°C. To detect PN-specific IgE, plates were incubated with biotinylated-CPE (biot-CPE; 0.1518 µg/mL in blocking solution) for 90 min at RT. Biot-CPE was generated using the EZ-Link Sulfo-NHS-LC-Biotinylation kit (Thermo Scientific). A standard curve, to which no capture antibody or sera was added, was generated using biot-CPE, serially diluted from 75 ng/mL to 0.6 ng/mL in PBS. Plates were washed thoroughly, then incubated with streptavidin-HRP (1:250 dilution in blocking solution; Thermo Scientific) and tetramethylbenzidine substrate (TMB; Thermo Scientific or Biolegend). The reaction was stopped with 1 N H_2_SO_4_. OD was measured at 450 nm, and the values were interpolated against the biot-CPE standard curve to determine relative PN-specific IgE-binding. Results are expressed as PN-IgE (Relative Binding, OD), reflecting the ability of serum PN-specific IgE to bind PN allergens.

### mMCP-1 quantification

Peripheral blood was collected by retro-orbital bleeding 40 min after the PN-challenge. The concentration of mMCP-1 was quantified using a Mouse mMCPT-1 ELISA kit (Thermo Scientific) following the manufacturer’s instructions. Briefly, serum samples were diluted by 1/25 and incubated on the capture antibody (anti-mouse mMCPT-1)-coated plate for 2 h at RT. After washing, a biotin-conjugated anti-mouse mMCPT-1 antibody was added to the plate and incubated for 1 h at RT. The reaction was developed with avidin-HRP and TMB, stopped with 1N H_2_SO_4_ and OD was measured at 450 nm. The results are expressed as ng of mMCP-1 per mL of serum and multiplied by the dilution factor of the sample.

### ELISA detection of PN allergens in biological samples

Ara h 1 and 2 were detected in samples harvested 40 min following intra-gastric administration of 24 mg of CPE in 0.8 mL of PBS. Serum from peripheral blood was collected by retro-orbital bleeding and analyzed undiluted. Whole small and large intestinal content was analyzed after dilution by 1/5 and 1/10 in PBS. The quantification of PN allergens was performed using proprietary ELISA kits for Ara h 1 and 2 following the manufacturer’s instructions (Indoor Biotechnologies). Briefly, for Ara h 1 detection by ELISA (Indoor Biotechnologies): Plates pre-coated with a monoclonal anti-Ara h 1 antibody were washed with wash buffer, and biological samples and Ara h 1 standards were added and incubated for 1 h. The plates were washed, and a second monoclonal anti-Ara h 1 antibody conjugated to peroxidase was added and incubated for 1 h. After a final wash, TMB substrate was added, and the reaction was stopped with 0.5 N H_2_SO_4_. OD was measured at 450 nm, and results were expressed as the concentration of Ara h 1 in biological samples, multiplied by the dilution factor. For Ara h 2 detection by ELISA (Indoor Biotechnologies): Plates pre-coated with a monoclonal anti-Ara h 2 antibody were washed with wash buffer, and biological samples and Ara h 2 standards were added and incubated for 1 h. After washing, a rabbit anti-Ara h 2 polyclonal detection antibody was added and incubated for 1 h. This was followed by the addition of a peroxidase-conjugated anti-rabbit IgG for 1 h. After the final wash, TMB substrate was added, and the reaction was stopped with 0.5 N H_2_SO_4_. OD was measured at 450 nm, and results were expressed as the concentration of Ara h 2 in biological samples, multiplied by the dilution factor.

### Detection of PN degradation by microbes isolated from human saliva and mouse and human intestinal content

Whole human saliva samples from healthy individuals (n=13), human small intestinal aspirates (jejunum) (n=5), and mouse small and large intestinal contents (10 mg/mL in PBS) were plated on agar media plates designed to isolate PN-degrading bacteria. BHI media enriched with powdered PN (organic partially defatted PN; PB&Me, Sahah Naturals) or agar supplemented with powered PN were used. Plates were incubated with samples for 48 h in aerobic and anaerobic conditions (Bactron IV anaerobic chamber). The bacteria were isolated based on their capacity to generate a visible hydrolytic halo in the media, which is indicative of the breakdown of PN components in the media. Isolated bacteria demonstrating PN hydrolytic activity were incubated in liquid BHI with CPE (0.5 mg/mL). After 48 h of incubation under aerobic or anaerobic conditions, the amount of Ara h 1 and 2 in media was quantified by ELISA as described above. Allergen degradation was calculated as the percentage of allergen degraded relative to the initial concentration in non-growth controls (CPE). These values were visualized in heatmaps using a blue-red color scale, where blue represents no degradation (0%) and red indicates complete degradation (100%).

Relative abundance of each bacterial isolate was determined by calculating the proportion of times each bacterial species was recovered among all PN-degrading isolates across all samples. Values were expressed as percentages and displayed using a green color scale, ranging from lightest green (lowest isolation frequency, minimum) to darkest green (highest isolation frequency, maximum).

Salivary and intestinal microbes were identified using Sanger sequencing technology. Briefly, DNA from isolates was extracted by picking single bacterial colonies into water, boiling, and centrifuging at 2,000 RCF for 2 min. The 8F-926R region of the 16S rRNA gene was amplified by PCR using the extracted DNA and sequences were determined using Sanger sequencing (FW primer: 5’-AGAGTTTGATCCTGGCTCAG-3’, RV primer: 5’-CCGTCAATTCCT-TTRAGTTT-3’). The resulting sequences for the isolates were taxonomically assigned using the NCBI nucleotide collection database^110^.

### PAF-AH quantification

Serum levels of platelet-activating factor acetylhydrolase (PAF-AH) were quantified using a mouse PAF-AH/PLA2G7/Lp-PLA2 ELISA kit (Novus Biologicals) according to the manufacturer’s instructions. Briefly, serum samples were diluted 1:10 in PBS and incubated capture antibody coated plates. After washing, a biotin-conjugated detection antibody was added for 1 h at RT, followed by HRP conjugate for 30 min. The reaction was developed with TMB, stopped with 1N H_2_SO_4_, and OD was measured at 450nm. Results were expressed as ng/mL serum and corrected for the dilution factor.

### Quantification of mucosal tryptase-positive mast cells

Paraffin-embedded small intestinal sections were deparaffinized in xylene and redhydrated through a graded ethanol series. Antigen retrieval was performed by heat in citric acid buffer, and non-specific binding was blocked with 3% BSA in PBS for 45 min. Sections were incubated o/n at 4°C with a rabbit monoclonal anti-mast cell tryptase antibody (1:200, Abcam). Immunoreactive cells were visualized using 3,3-diaminobenzidiene (DAB; Dako). Negative controls were performed by omitting the primary antibody. Tryptase positive mast cells were quantified in the mucosa and normalized to mucosal area using an Axio Imager Z2 microscope (Carl Zeiss, Jena, Germany). The data are expressed as tryptase-positive cells/µm^2^.

### Microbial digestion of PN for challenge

Selected bacteria (*Rothia mucilaginosa* (R3), *Staphylococcus epidermidis* (S1), and *Staphylococcus aureus* (S2)) were grown o/n on BHI agar plates enriched with 1% powdered PN to confirm PN degradation capability (visible PN hydrolytic halo). For each strain, a 10 mL pre-inoculum was prepared by o/n culture in liquid BHI. Then, 10 µL of pre-inoculum was transferred into 5 mL of OptiMEM medium containing 30 mg/mL CPE and incubated for 6 h to complete the PN digestion process. The resulting digests were boiled for 5 min to kill the bacteria, and 800 µL of the digests, containing 24 mg of digested PN, was administered by oral gavage at the time of challenge as described above. OptiMEM medium served as the vehicle for non-digested CPE challenge.

### Measurement of *ex vivo* intestinal PN-degrading capacity

The ability of the intestinal content from mice colonized with PN-degrading bacteria to degrade PN was assessed using ELISA and a bioassay with agar media enriched with PN. Solid BHI agar enriched with 1% powdered PN was prepared, and a 50 μL aliquot of diluted (1/2 in PBS) small intestinal content was placed into wells created in the agar. The plates were then incubated for 24 h. The PN-degrading capacity was determined by measuring the diameter of the clear halo around the inoculation site, indicating degradation, and the results were expressed in millimeters. In addition, the intestinal contents were also incubated with CPE (0.5 mg/mL). After 48 h of incubation, the quantity of Ara h 1 and 2 in media was analyzed by ELISA as described above. The data for Ara h 1 and 2 degradation are represented as blue-red color scales showing the percentage of allergen degradation relative to the control incubations (CPE), with blue indicating no degradation (0%) and red indicating complete degradation (100%).

### Ussing chamber detection of PN translocation after bacterial digestion

Translocation of PN allergens after digestion by bacteria was evaluated *ex vivo* using small intestinal sections of SPF C3H/HeN mice and the Ussing chamber technique. For preparation of bacterial PN allergen digestions, *Rothia mucilaginosa* (R3), *Staphylococcus epidermidis* (S1), and *Staphylococcus aureus* (S2 & S3) were grown for 16 h in BHI liquid medium. Bacteria were then incubated in BHI with 5 mg/mL CPE for 4 h to allow for bacterial digestion of the PN allergens. After incubation, samples were boiled for 15 min. Bacterial digestions were performed in triplicate, with a negative control containing no bacterial inoculum. For the Ussing chamber technique, 1.5 cm sections of proximal small intestine were collected, opened along the mesenteric border, and mounted on the sliders of Ussing chambers. Each chamber exposed 0.25 cm² of tissue surface area to 4 mL of circulating oxygenated Krebs buffer containing 10 mM glucose (serosal side) and 10 mM mannitol (mucosal side), maintained at 37°C and aerated with 95% O_2_ and 5% CO_2_. Then, 0.2 mL of each bacterial digestion was placed into 4 mL of Krebs buffer on the mucosal side of the chamber. The experiment was run for 2 h, after which the total mucosal and serosal volumes were collected. The concentrations of PN allergens Ara h 1 and Ara h 2 on the serosal side were quantified by ELISA and expressed as intestinal translocation of allergens. The Ussing chamber system was from Physiologic Instruments.

### Microbial digestion of PN for proteomic and immune activation assays

*Rothia* (R1–*Rothia aeria* R2–*Rothia dentacariosa* R3–*Rothia mucilaginosa*) and *Staphylococcus* (S1–*Staphylococcus epidermidis,* S2–*Staphylococcus aureus,* S3–*Staphylococcus aureus*) species were selected for characterization of PN allergen degradation and allergenicity. Bacteria were grown o/n in BHI (10 mL), then 10 µL of bacterial culture was taken and used to inoculate 1 mL of OptiMem with 0.5 mg/mL CPE. The bacteria were incubated in the media with CPE for 4 h at 37°C. After the incubation, the digestion was stopped by boiling for 5 min. Subsequently, the digestion of Ara h 1 and 2 were detected using ELISA, SDS-PAGE and Western blotting, and BMMC activation assays.

### SDS-PAGE and Western blotting

To assess PN degradation profiles produced by degradation by *Rothia* and *Staphylococcus* species we performed SDS-PAGE and Western blotting. Bacteria were grown in 0.5 mg/mL as mentioned above. Undigested PN and Ara h 1 and 2 were used as controls. Then Laemmli sample buffer (Bio-Rad) with β-mercaptoethanol (Sigma-Aldrich) at 1/10 was added at 1/4 in a final volume of 50 µL. Before loading the gel (Any kD™ Mini-PROTEAN® TGX™ Precast Protein Gels, Bio Rad), samples were heat-denatured at 100°C for 5 min. Moreover, a protein molecular weight marker was added (ThermoFischer). The gel was run in 1X running buffer (*i.e.*, SDS/Tris/Glycine 10X, Bio-Rad) for 1-1.5 h at 100 V using an electrophoretic cell and a power supply Mini-Protean System Bio-Rad (Bio-Rad).

For SDS-PAGE, after running, the gel was placed in staining solution (45% of methanol and 10% of acetic acid in distilled water, plus 0.2% w/v of Coomassie blue, Bioscience) for 1 h in agitation. Then, the staining solution was removed using destaining solution (20% methanol and 10% acetic acid, Sigma-Aldrich, in distilled water) in constant agitation, the process was repeated every 15 min until the background was eliminated. Then, the gel was photographed using a molecular imaging system (Amersham ImageQuant™ 800).

For Western blotting, after running the gel, the bands of the gel were transferred to a nitrocellulose membrane (Bio-Rad) using a transfer system (fiber pad, filter paper, membrane, gel, filter paper and fiber pad) and transfer buffer (20% methanol and 10% of Tris-Glycine 1X in distilled water). The transference was run for 1 h at 400 mA. Then, to check that bands were transferred correctly, the membrane was stained with Ponceau S Staining solution (ThermoScientific). After cleaning with water and 3 washes with TBS-T solution, the membrane was blocked in constant agitation for at least 1 h with 3% BSA (NZYtech) in TBS-T. Then, the membrane was incubated with sera from PN-allergic donors, or sera from mice allergic to Ara h 1 or to Ara h 2 (diluted in 3% BSA at different dilutions depending on the amount of IgE) as primary antibody, at 4°C o/n with smooth agitation. Sera from non PN-allergic donors (Table S2) and non PN-allergic mice were used as negative control. During day 2, after performing 6 washes with TBS-T every 10 min with agitation, the membrane was incubated with anti-human IgE-HRP (Invitrogen) 1:10,000 or anti-mouse IgE-HRP (SouthernBiotech) 1:1,000 as secondary antibody for 1 h at RT with agitation. Then, 6 washes with TBS-T every 10 min with agitation were performed before revealing the membrane According to the instructions of the manufacturer (Cytiva-Merck), 1:1 of peroxide and luminol and 1 mL of the mixture was added to the membrane. The chemiluminescent signal of the gel was captured in a chemiluminescent imaging system (Amersham ImageQuant™ 800). SDS-PAGE and Western blots were analyzed using Fiji ImageJ.

### Proteomics

Proteomics was performed to identify bacterial degradation products of PN allergens and to map PN-specific epitopes. Microbial PN digestions (above) by *Rothia* (R1, R2, and R3) and *Staphyloccocus* (S1, S2, and S3) were ultra-filtered using microcon filter units with a 3 kD cutoff that were previously equilibrated and washed with water. The peptides of the flow-through were selected (<3 kD) and they were cleaned with ZipTip with 0.6 µL C18 resin (Merck Millipore) to avoid the presence of elements that could interfere with the mass spectrometry, and they were drained in a SpeedVac Vacuum Concentrator. The samples, resuspended in a volume of 14 µL and 2 µL LC-MS grade water containing 2% (v/v) acetonitrile and 0.1% (v/v) formic acid, were used for the quantification with the QBIT method, analyzing around 0.8 µg per sample. The peptides were analyzed by nano liquid chromatography coupled to mass spectrometry in data-dependent-acquisition mode using an UltiMate 3000 High Pressure Liquid Chromatograph (Fisher Scientific) and a Orbitrap Exploris™ 240 Mass Spectrometer (Fisher Scientific). The flow of the chromatograph was 250 nL/min for 60 min, and an Easy-spray PepMap C18 analytical column 50 cm × 75 μm (Fisher Scientific) was used.

The obtained MS1 and MS2 spectra were analyzed using the Peaks (https://www.bioinfor.com/peaksdb/) search engine against an *Arachis hypogaea* proteome database obtained from UniprotKB (https://www.uniprot.org/) combined with a list of typical laboratory contaminants, using a tolerance of 10 ppm and 0.02 Da for the precursor ions and fragments, respectively. Carbamidomethylation in cysteines was used as fixed modification and acetyl in the N-terminal end of the protein and methionine oxidation were used as variable modifications. The search was performed without restrictions to any proteases and the results were shown as proteins identified with at least a unique peptide with a False Discovery Rate ≤1%.

The unique peptides obtained after the analysis were located and identified in the sequences of Ara h 1, which were downloaded from the Protein Data Bank (https://www.rcsb.org/). The most common human IgE-binding epitopes of Ara h 1 were found in the following allergen databases: Allergen Nomenclature (https://allergen.org/); Allergen Online (http://allergenonline.org/); Compare Database (https://db.comparedatabase.org/); and The Immune Epitope Database ( https://www.iedb.org/). The following studies were considered to specify immunodominant IgE-binding epitopes^35-46^. Ara h 1 peptides that were contiguous or overlapping were considered distinct if their cut-off points differed. Peptides not fully aligned with the canonical sequence were required to have at least 75% alignment and a minimum of four consecutive identical amino acids Molecular structure analysis was performed with Chimera v1.17.3 for tridimensional visualization of allergens and relevant epitopes before and after bacterial digestion.

### Mice sensitization to Ara h 1 and Ara h 2

Mice were sensitized to immunodominant PN allergens to generate sera specific to Ara h 1 or Ara h 2 specific IgE to be used in mast cell degranulation assays (above). To sensitize mice against native Ara h 1 (InBio) or recombinant Ara h 2 (prepared as reported^111^ using Uniprot reference Q6PSU2-1 without the signal peptide), 3 i.p. injections with 10 µg of the allergen plus 1 mg aluminum hydroxide (Alhydrogel® adjuvant 2%, InvivoGen) were performed. Previously, aluminum hydroxide and allergen were mixed at 4°C during 30 min in a ferris wheel. Between the first and second injection, 2 weeks were left; and 1 week between the second and third injection. Blood samples were collected and centrifuged at 16,000 RCF to obtain sera for mast cell activation assays (see below) and Western blot analysis. Control sera from PN-allergic mice were generated following immunization with a classical model of food allergy.^48,72^

### Bone marrow-derived mast cell culture

BMMCs were cultured to assess mast cell activation by bacterially digested PN allergens. Bone marrow from femur and tibia of C57BL/6 (Charles River) mice were used to obtain BMMCs as described^112^. In short, bone marrow was flushed out inserting a 23-gauge needle (BD Microlance) attached to the 10 mL syringe filled with PBS (Gibco) at the knee side of both types of bone. Then, the cell suspension collected was filtered with a 40 µm filter (Falcon). After centrifugating 5 min at 286 RCF and RT, the pellet was lysed with ACK lysing buffer (Lonza). Erythrocyte-lysed bone marrow cells were cultured for 4 weeks in Petri dishes at 37°C and 5% CO_2_ in Iscove’s Modified Dulbeccós Medium (IMDM, Gibco) supplemented with 10% fetal bovine serum (FBS, Cytiva), 1% minimal essential medium (MEM, Gibco), 1% sodium pyruvate (Biowest), 1% penicillin/streptomycin (P/S, Gibco), 1% non-essential amino acids (Biowest), 0.01% recombinant murine stem cell factor (SCF, PeproTech) and 0.05% recombinant murine Interleukin (IL-3, PeproTech), until they differentiated into mature BMMCs. Cell growth and viability were monitored using the trypan blue exclusion test. The culture media was changed weekly, and the cellularity was kept below 0.5 × 10^6^ cells/mL.

### BMMC activation

BMMC activation was conducted as reported^47,48^. In brief, BMMCs were sensitized o/n at 37°C and 5% CO_2_ with sera at a concentration range of 15–40 ng/mL from Ara h 1- or Ara h 2-sensitized mice. Culture media was changed 1-2 days previous sensitization. PN-allergic mice sera were used as a positive control. BMMC sensitization was done at a cell density of 1 × 10^6^ cells/mL. The next day, BMMCs were washed to eliminate unbound Igs (cells were spun at 265 RCF for 5 min and low break) and resuspended in supplemented IMDM without cytokines, nor FBS, at a cell density of 1 × 10^6^ cells/mL. For BMMC activation, bacteria with PN-degrading potential isolated from saliva and gut of healthy donors were selected. Then, 0.1 × 10^6^ BMMCs per condition were placed in a U/V bottom 96-well culture plate (Falcon) and challenged with bacterially digested PN (Stallergenes Greer) at 25 µg/mL and 100 µg/mL in Ara h 1-sensitized BMMCs, or at 50 µg/mL and 200 µg/mL in Ara h 2-sensitized BMMCs. Media and undigested PN were used as negative and positive controls, respectively. After 20 min at 37°C and 5% CO_2_, the reaction was stopped on ice and BMMC activation was determined via β-hexosaminidase activity in the supernatants, and phenotypically (CD63 and CD107a expression) by flow cytometry^47^.

### β-hexosaminidase activity assay

BMMCs were centrifugated as above described to recover cell-free supernatants. A volume of supernatant of 45 µL was added in duplicates to 45 µL of β-hexosaminidase substrate solution (2 mM p nitrophenyl N-acetyl β-D-glucosamine, Sigma-Aldrich) diluted in 0.1 M citrate buffer (45 mM dehydrated sodium citrate, 55 Mm citric acid, Sigma-Aldrich, in distilled H_2_O) in a flat-bottom 96-well plate. After incubation at 37°C for 45 min in the dark, 45 µL NaOH 1 M (1M sodium hydroxide, Sigma-Aldrich, in distilled H_2_O) were added to stop the reaction. OD was measured at 405 nm with a microplate reader. Buffers alone and BMMC lysates (0.5% Triton X-100, Sigma Aldrich, was used for cell lysis) were used as control, absorbance background (B) and total lysis, respectively. The percentage of BMMC degranulation, based on β-hexosaminidase activity, was calculated as follows:

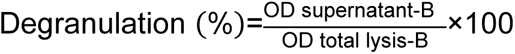

### Flow cytometry

BMMCs were resuspended in ice-cold FACS buffer (2.5 mM EDTA, PanReacAppliChem, 0.5% BSA in PBS). BMMCs were blocked with 1:50 Fc block (purified anti-mouse CD16/32, BioLegend) for 15 min on ice to prevent non-specific antibody binding. Then, BMMCs were incubated with BV421 rat anti-CD117 (c-kit) (BioLegend) 1/200; APC rat anti-CD63 (BioLegend) 1:200; PerCP-Cy5.5 rat anti-CD107a (LAMP-1) (BioLegend) 1:200; PE rat anti-IgE (BioLegend) 1:200; PE-Cy7 rat anti-FcεR1α (BioLegend) 1:200; on ice for 30 min covered from light. Viability was assessed with efluor780 dye (eBioscience) 1:4,000. After washing with FACS buffer to eliminate unbound antibodies, cells were analyzed on a BD FACSCantoll flow cytometer. On average, 10,000 live and single cells were recorded. Data were analyzed with FlowJo v10 software; dead cells and aggregates were excluded and fluorescence minus one (FMO) controls were used for gating.

### Microbiota analysis

DNA was extracted from mouse fecal pellets, human saliva, and human jejunal aspirates, and the hypervariable V3-V4 regions of the 16S rRNA gene were amplified with polymerase chain reaction (PCR) using Taq polymerase (Life Technologies), as previously described^82^. Forward barcoded primers targeting the V3 region (v3f_341f-CCTACGGGNGGCWGCAG) and reverse primers targeting the V4 region (v4r_806r-GGACTACNVGGGTWTCTAAT) were used. Forward primers included six-base pair barcodes to allow multiplexing samples. Purified PCR products were sequenced using the Illumina MiSeq platform by the McMaster Genomics facility. Primers were trimmed from the obtained sequences with Cutadapt software^113,114^, and processed with Divisive Amplicon Denoising Algorithm 2 (DADA2; version 1.14.0) using the trained SILVA reference database (version 138.1)^115,116^. A phylogenetic tree of the sequences was calculated using FastTree 2^117^, and data was explored using the phyloseq package (version 1.30.0) in R (version 3.6.2)^118^. After data cleanup, a total of 292,618 reads were obtained with a minimum of 11,651 and maximum of 60,790 with an average of 32,513 reads per sample for mouse microbiota. For human healthy control salivary microbiota, a total of 273,686 reads were obtained with a minimum of 1,881 and maximum of 48,620 with an average of 16,099 reads per sample. For PN allergic patient salivary microbiota, a total of 844,167 reads were obtained with a minimum of 3,236 and maximum of 145,324 with an average of 36,702 reads per sample. Alpha-diversity was measured using observed species and Chao1 indices. Beta-diversity was calculated on normalized data and the originated matrices were ordinated using principal coordinate analysis based on Jaccard distance (mouse microbiota) and Bray-Curtis dissimilarity (human microbiota).

For whole genome sequencing of bacterial isolates, genomic DNA was extracted as previously described^82^. Illumina short-read libraries were prepared using the NEBNext Ultra II FS DNA Library Prep Kit for Illumina (NEB, Ipswich, MA, USA). Final libraries were sequenced on an Illumina NextSeq 2000 platform using a P1 flow cell at the McMaster Metagenomics Facility (Hamilton, ON, Canada). The run yielded 3.7 million paired-end (2 × 150 bp) reads per genome. Short reads were processed as previously described^119^, with few modifications. Briefly, fastp was used for quality trimming and removal of adapters from Illumina reads^120,121^. Unicycler (v.0.5.1) was then used for de novo assembly^122^, followed by Bakta for annotation of the genomes^123^. Genomes were then compared using Roary^124^. The data was further analysed using the MEROPs database and R (4.4.3) to search for proteases. The data are presented as heatmaps where light blue=gene not present and dark blue=present.

*Rothia* abundance (16S rRNA) data in saliva of non-PN allergic and PN-allergic (high threshold and low threshold) individuals were obtained from a previously published PN allergy cohort described by Zhang *et al*^52^. PN threshold was determined as described in the primary study^52^. In this cohort, amplicon sequence variants (ASVs) belonging to *Rothia* were identified after filtering for low prevalence and low abundance, and relative abundances were quantified following taxonomic assignment against NCBI databases. We used the *Rothia* ASV abundance values as provided by the authors for secondary analysis in the present study.

## STATISTICAL ANALYSIS

All variables were analyzed using GraphPad Prism 9 and 10 software (GraphPad Software, USA). Parametric data are depicted as dot plots with each dot representing an individual mouse or biological replicate. Normal distribution was determined by D’Agostino-Pearson omnibus normality test, Shapiro-Wilk test, and Kolmogorov-Smirnov test with Dallal-Wilkinson-Lillie correction. One way analysis of variance (ANOVA) was used to evaluate differences between more than two groups with a parametric distribution and Tukey’s or Dunnet’s post-hoc corrections were applied. Student’s *t*-test (two-tailed) was performed to evaluate the differences between two independent groups as appropriate. Data with non-normal distribution were evaluated with Kruskal-Wallis test with Dunn’s post-hoc test for more than two groups. Correlation analyses were performed using Pearson correlation for parametric data and Spearman correlation for non-parametric data. A *P* value of ≤ 0.05 was selected to reject the null hypothesis. Information regarding specific *P* values, value of *n*, and how data are presented can be found in figure legends.

