## Supplementary Figures and Tables for "Microbial metabolism of food allergens determines the severity of IgE-mediated anaphylaxis"

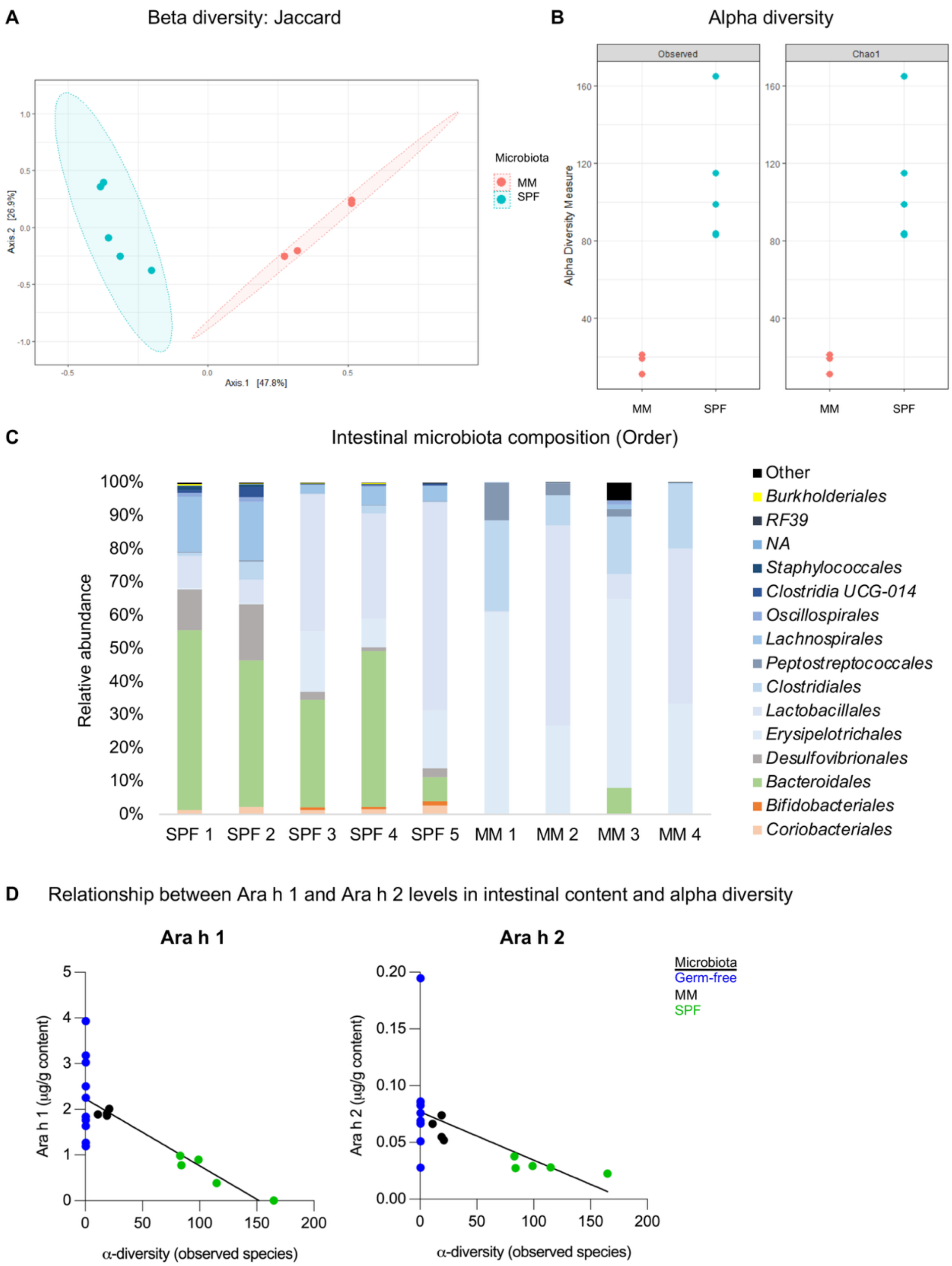


**Figure S1.** Intestinal microbiota of minimal microbiota (MM) and specific pathogen-free (SPF) C57BL/6 mice. (A) Beta-diversity (Jaccard index) plot of fecal microbiota where each dot represents one mouse. (B) Alpha-diversity metrics based on observed species and Chao1 index where each dot represents one mouse. (C) Order-level relative abundance of the microbial composition where each bar represents one mouse. (D) Correlation plot of Ara h 1 ($y$=-0.01467x + 2.231, $R^2$=0.5678, $P$=0.0002) and Ara h 2 ($y$=-0.0004257x + 0.07693, $R^2$=0.3076, $P$=0.0137) against alpha-diversity in small intestinal content of germ-free, MM, and SPF mice. n=4–5 mice per group from one experiment (A–C).

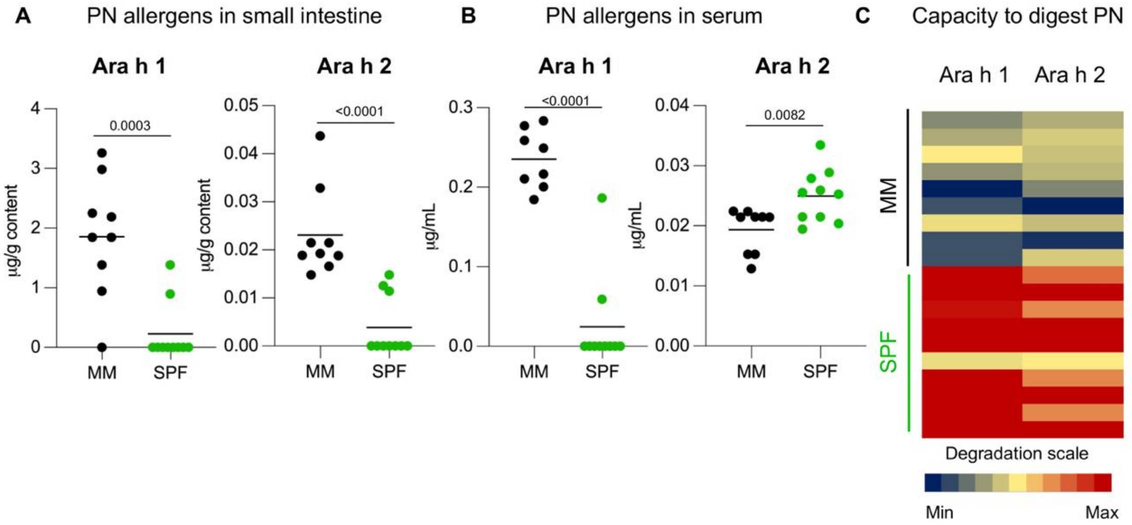


**Figure S2.** Peanut (PN) digestion by minimal microbiota (MM) and specific pathogen-free (SPF) C3H/HeN mice. Mice were provided a 24 mg bolus of PN i.g. Peripheral blood and small intestinal contents were collected 40 minutes post PN delivery for analysis. PN allergens (Ara h 1 and 2) in small intestinal content (A) and in serum from peripheral blood (B). (C) Heatmap showing digestion capacity of the microbiota against Ara h 1 and 2. Small intestinal contents were incubated with PN allergens *in vitro* and remaining allergens were quantified after digestion. Allergen degradation capacity is expressed as the percentage of degraded allergen relative to control non-growth incubations (containing crude peanut extract, CPE), with the color scale ranging from blue (0% degradation; minimum) to red (100% degradation; maximum). Significance levels: Ara h 1 SPF vs. MM ($P$<0.0001), Ara h 2 SPF vs. MM ($P$<0.0001). n=8–10 mice per group; pooled from 2 independent experiments. Data are presented as mean where each dot represents an individual mouse (A–B), or each row represents one mouse (C). Displayed $P$ values were calculated using an unpaired Student's $t$-test.

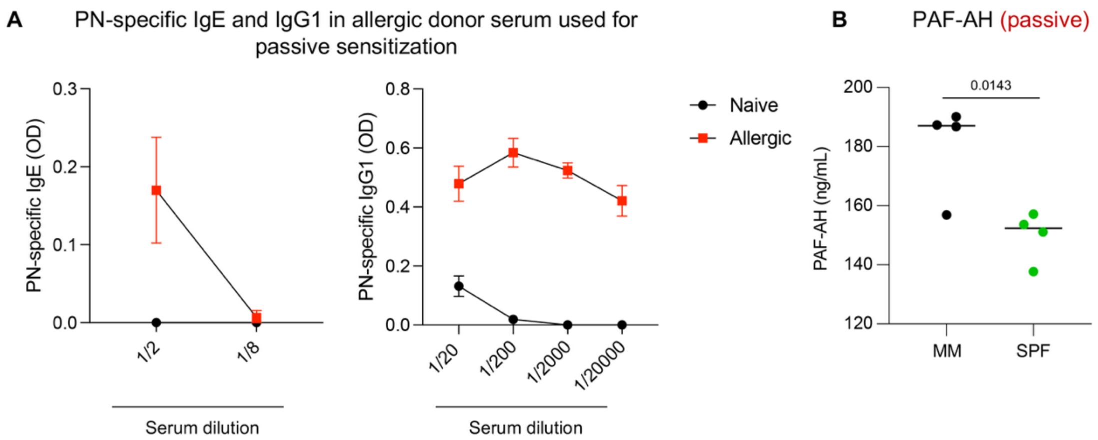


**Figure S3.** (A) PN-specific IgE and IgG1 in allergic donor serum used for passive sensitization (n=4). (B) Serum platelet-activating factor acetylhydrolase (PAF-AH) after peanut (PN)-challenge for passively sensitized mice (n=4 from one experiment). (B). Data are presented as mean ± SD (A) and mean where each dot represents one mouse (B). Displayed *P* values were calculated using an unpaired Student's *t*-test.

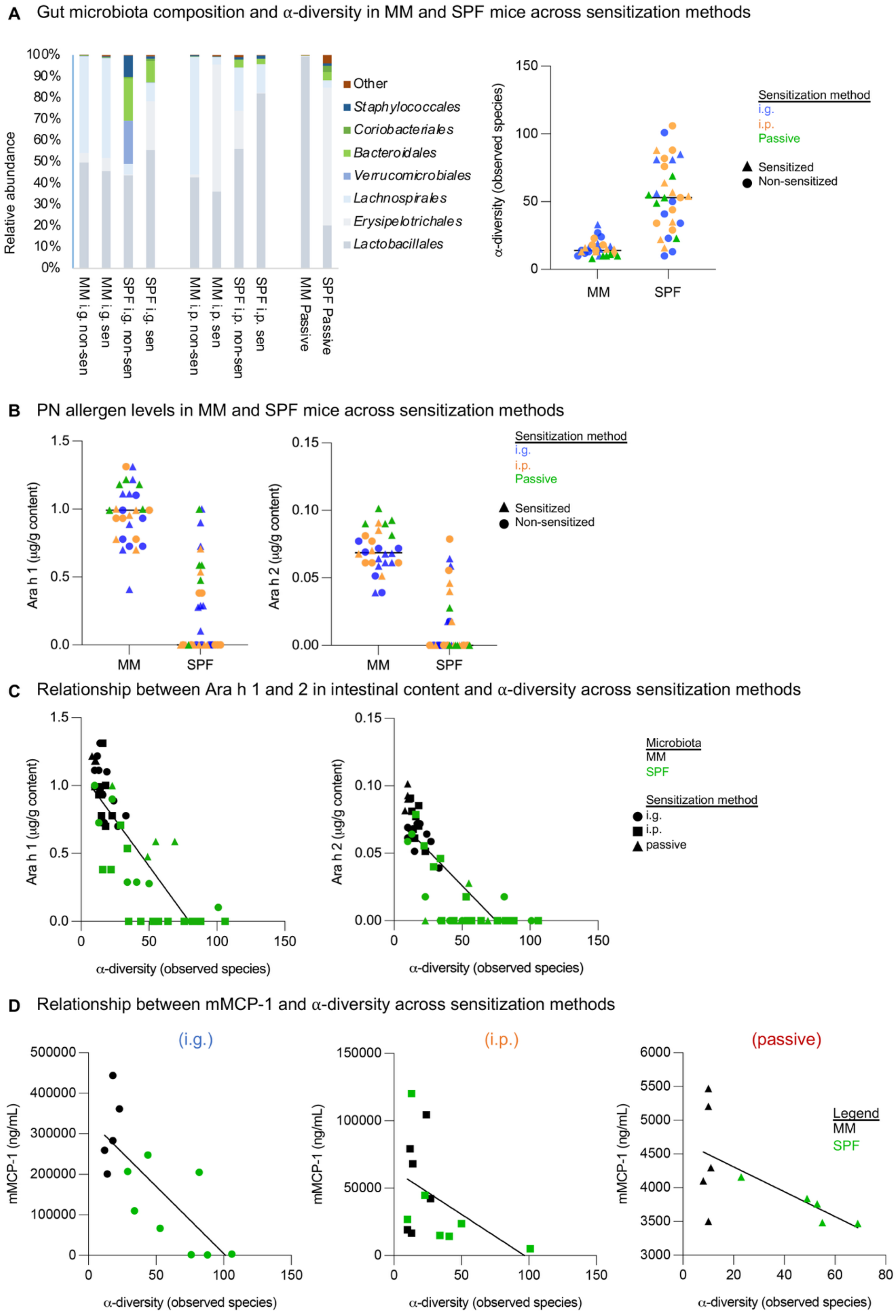
A Gut microbiota composition and α-diversity in MM and SPF mice across sensitization methods
Relative abundance
100%
90%
80%
70%
60%
50%
40%
30%
20%
10%
0%
MM i.g. non-sen
MM i.g. sen
SPF i.g. non-sen
SPF i.g. sen
MM i.p. non-sen
MM i.p. sen
SPF i.p. non-sen
SPF i.p. sen
MM Passive
SPF Passive
Other
Staphylococcales
Coriobacteriales
Bacteroidales
Verrucomicrobiales
Lachnospirales
Erysipelotrichales
Lactobacillales
α-diversity (observed species)
150
100
50
0
MM
SPF
Sensitization method
i.g.
i.p.
Passive
Sensitized
Non-sensitized
B PN allergen levels in MM and SPF mice across sensitization methods
Ara h 1 (µg/g content)
1.5
1.0
0.5
0.0
MM
SPF
Ara h 2 (µg/g content)
0.15
0.10
0.05
0.00
MM
SPF
Sensitization method
i.g.
i.p.
Passive
Sensitized
Non-sensitized
C Relationship between Ara h 1 and 2 in intestinal content and α-diversity across sensitization methods
Ara h 1 (µg/g content)
1.5
1.0
0.5
0.0
0
50
100
150
α-diversity (observed species)
Ara h 2 (µg/g content)
0.15
0.10
0.05
0.00
0
50
100
150
α-diversity (observed species)
Microbiota
MM
SPF
Sensitization method
i.g.
i.p.
passive
D Relationship between mMCP-1 and α-diversity across sensitization methods
(i.g.)
mMCP-1 (ng/mL)
500000
400000
300000
200000
100000
0
0
50
100
150
α-diversity (observed species)
(i.p.)
mMCP-1 (ng/mL)
150000
100000
50000
0
0
50
100
150
α-diversity (observed species)
(passive)
mMCP-1 (ng/mL)
6000
5500
5000
4500
4000
3500
3000
0
20
40
60
80
α-diversity (observed species)
Legend
MM
SPF

**Figure S4.** Gut microbiota composition and diversity of minimal microbiota (MM) and specific pathogen-free (SPF) C3H/HeN mice across sensitization methods, and their relationship with peanut (PN)-challenge outcomes (corresponding to Figure 2). (A) Relative abundance of bacterial orders and α-diversity (observed species) in the small intestine across intragastric (i.g.), intraperitoneal (i.p.), and passive sensitization, including sensitized non-sensitized mice. (B) Levels of Ara h 1 and 2 in intestinal content across sensitization methods, including sensitized and non-sensitized mice. (C) Correlation plot of Ara h 1 ($y$=-0.01375x + 1.095, $R^2$=0.7053, $P$<0.0001) and Ara h 2 ($y$=-0.001039x + 0.07793, $R^2$=0.6736, $P$<0.0001) against α-diversity in small intestinal content. (D) Relationship between serum mucosal mast cell protease 1 (mMCP-1) concentrations and α-diversity across sensitization methods.

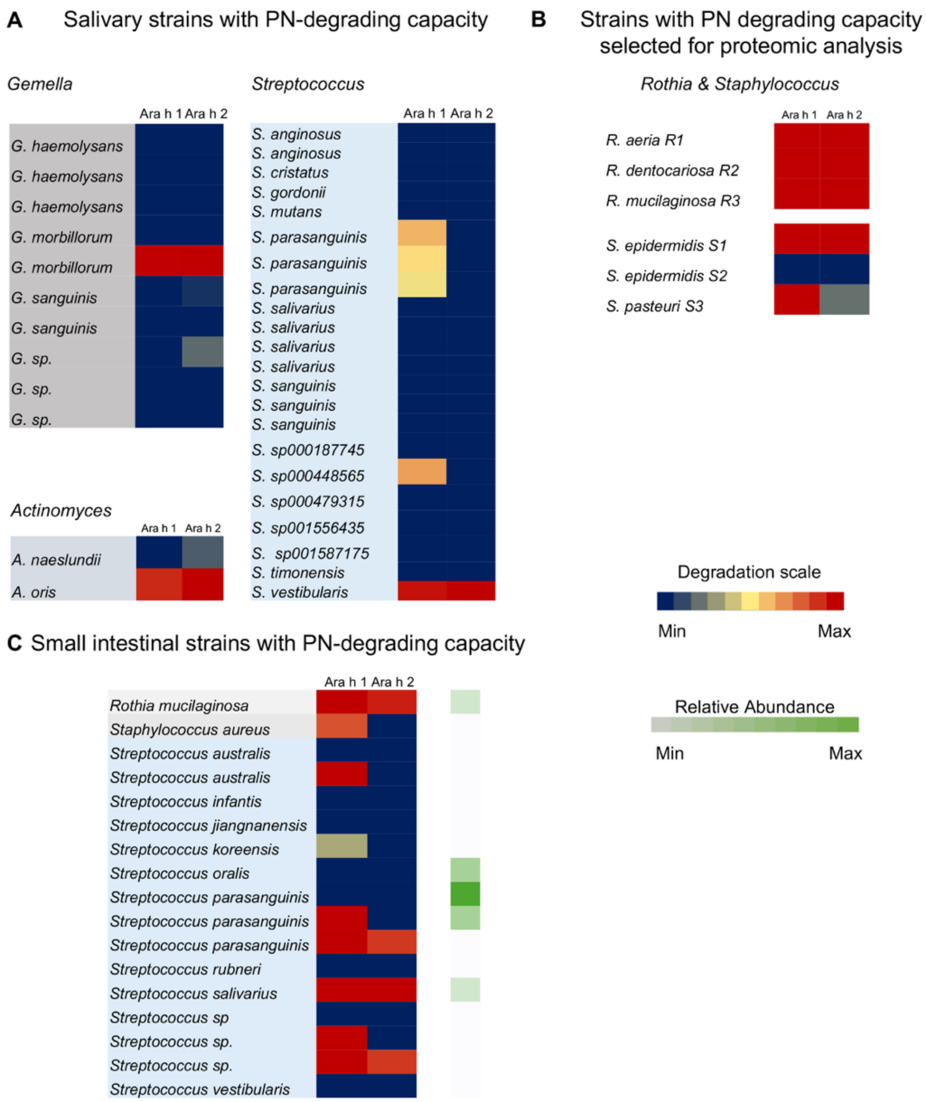


**Figure S5.** Characterization of peanut (PN)-degrading bacteria isolated from human saliva and small intestine. (A) Heatmap of PN allergen degradation capacity of bacterial strains isolated from saliva. (B) Degradation of Ara h 1 and 2 by bacterial strains that were further characterized in Figures 4 and 5 for their proteomic profiles and effects on mast cell activation. (C) Heatmap of PN allergen degradation capacity of bacterial strains isolated from the small intestine (jejunal aspirates). Color scale (blue-red) represents the percentage of allergen degradation relative to control non-growth incubations, with blue indicating no degradation and red indicating complete degradation. For (C), the relative abundance (green scale) reflects how often each species was isolated out of all PN-degrading isolates, with darker green indicating more frequently isolated species (light=low, dark=high).

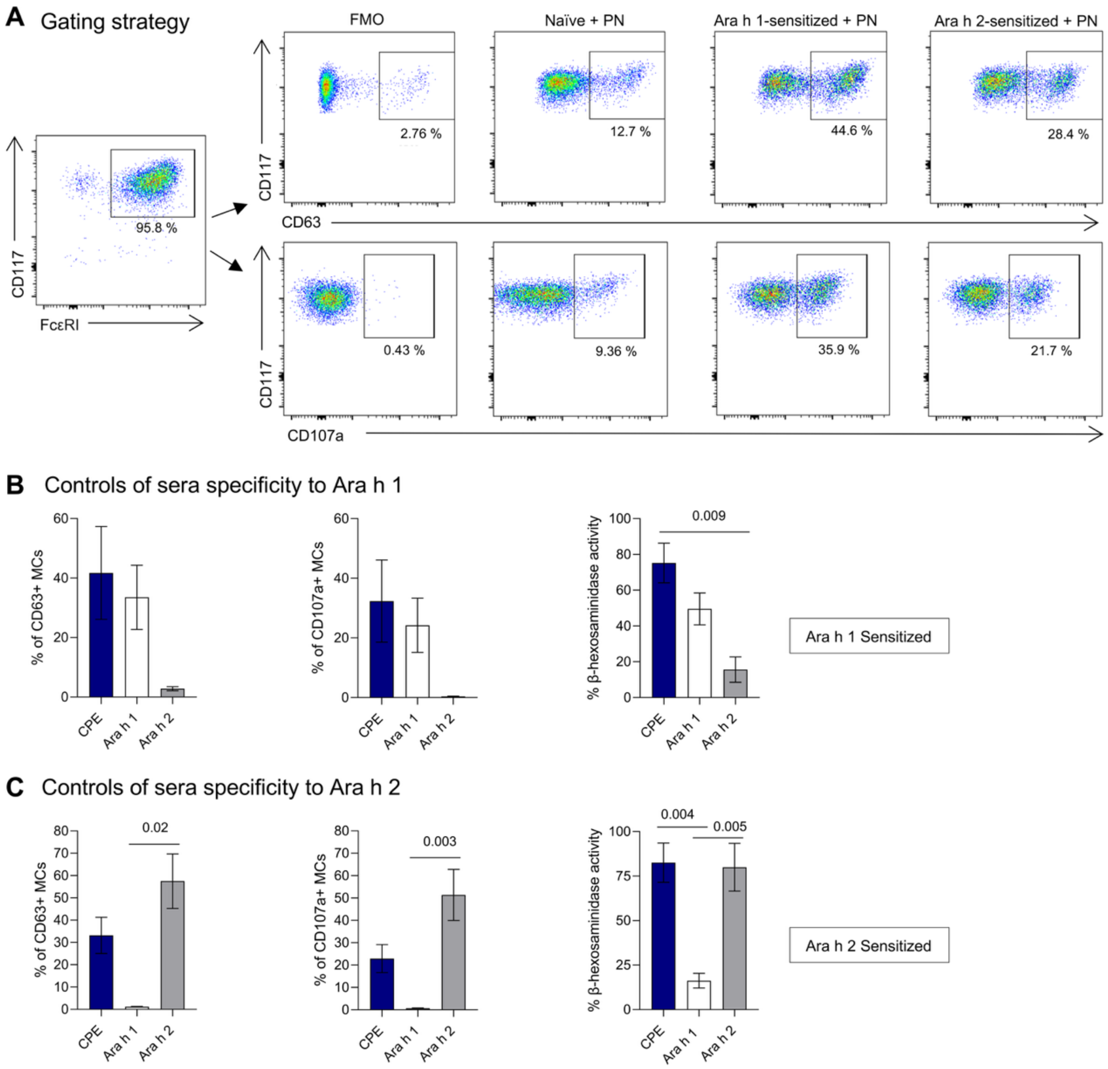


**Figure S6.** Characterization of sera specificity from mono-sensitized mice. (A) Gating strategy. Bone marrow-derived mast cells (MCs) co-express FcεRI and CD117. MC activation was detected by the expression of CD63 and CD107a. Gates of positive and negative controls are shown. Controls of sera specificity to (B) Ara h 1 and (C) Ara h 2. MCs were sensitized with a pool of sera from Ara h 1- or Ara h 2-allergic mice, respectively, and then challenged with undigested peanut (PN, blue), native Ara h 1 (white) or recombinant Ara h 2 (grey). Data are presented as the mean +/- SEM of each group. Displayed *P* values were calculated using a one-way ANOVA with Tukey's post-hoc test in panels B and C (CD107a and β-hexosaminidase activity) or using Kruskal-Wallis test with Dunn´s post hoc test in panel C (CD63).

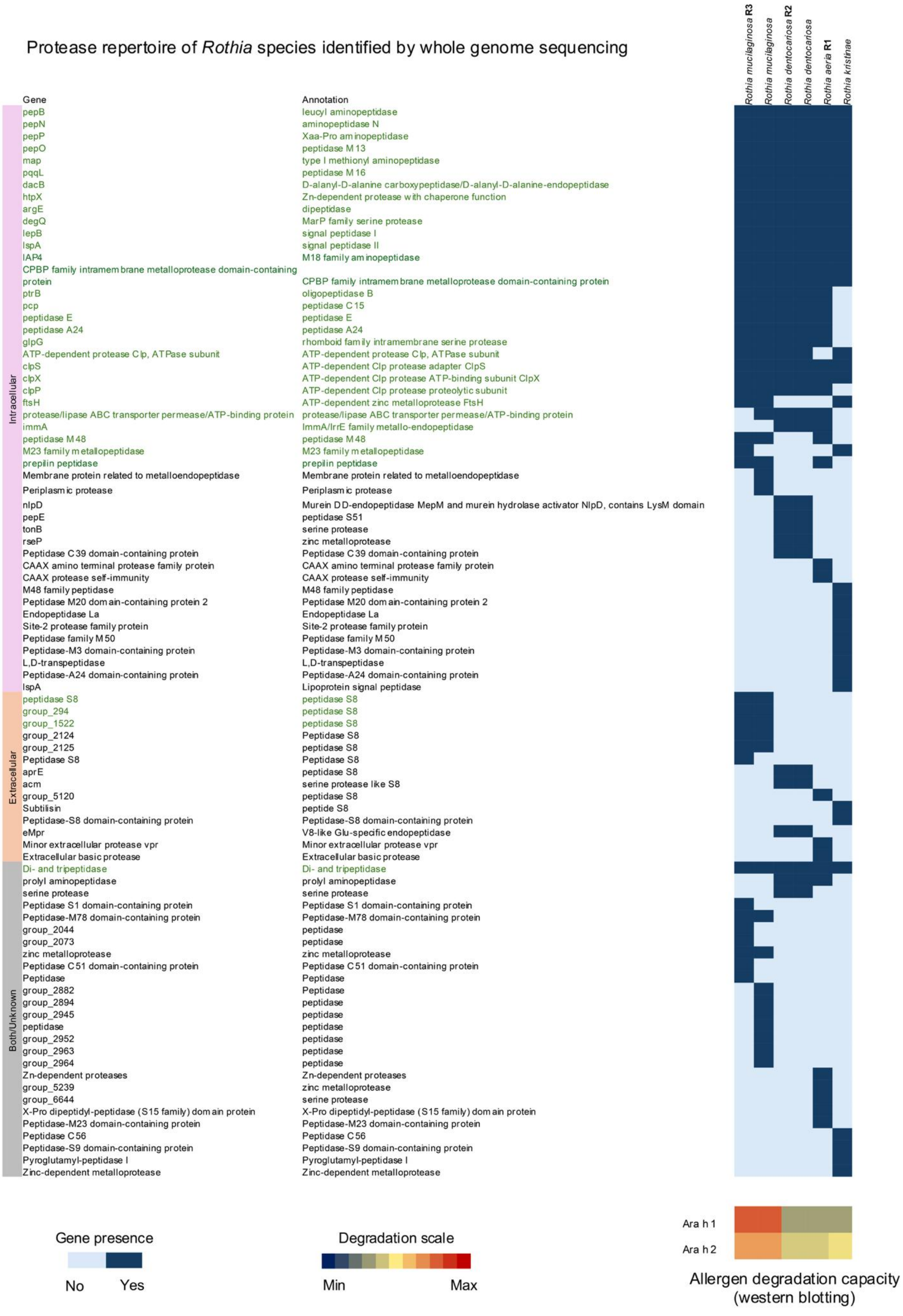
Protease repertoire of Rothia species identified by whole genome sequencing
Rothia mucilaginosa R3
Rothia mucilaginosa
Rothia dentocariosa R2
Rothia dentocariosa
Rothia aeria R1
Rothia kristinae
Gene
Annotation
Intracellular
pepB leucyl aminopeptidase
pepN aminopeptidase N
pepP Xaa-Pro aminopeptidase
pepO peptidase M13
map type I methionyl aminopeptidase
pqqL peptidase M16
dacB D-alanyl-D-alanine carboxypeptidase/D-alanyl-D-alanine-endopeptidase
htpX Zn-dependent protease with chaperone function
argE dipeptidase
degQ MarP family serine protease
lepB signal peptidase I
lspA signal peptidase II
IAP4 M18 family aminopeptidase
CPBP family intramembrane metalloprotease domain-containing protein CPBP family intramembrane metalloprotease domain-containing protein
ptrB oligopeptidase B
pcp peptidase C15
peptidase E peptidase E
peptidase A24 peptidase A24
glpG rhomboid family intramembrane serine protease
ATP-dependent protease Clp, ATPase subunit ATP-dependent protease Clp, ATPase subunit
clpS ATP-dependent Clp protease adapter ClpS
clpX ATP-dependent Clp protease ATP-binding subunit ClpX
clpP ATP-dependent Clp protease proteolytic subunit
ftsH ATP-dependent zinc metalloprotease FtsH
protease/lipase ABC transporter permease/ATP-binding protein protease/lipase ABC transporter permease/ATP-binding protein
immA ImmA/IrrE family metallo-endopeptidase
peptidase M48 peptidase M48
M23 family metallopeptidase M23 family metallopeptidase
prepilin peptidase prepilin peptidase
Membrane protein related to metalloendopeptidase Membrane protein related to metalloendopeptidase
Periplasmic protease Periplasmic protease
nlpD Murein DD-endopeptidase MepM and murein hydrolase activator NlpD, contains LysM domain
pepE peptidase S51
tonB serine protease
rseP zinc metalloprotease
Peptidase C39 domain-containing protein Peptidase C39 domain-containing protein
CAAX amino terminal protease family protein CAAX amino terminal protease family protein
CAAX protease self-immunity CAAX protease self-immunity
M48 family peptidase M48 family peptidase
Peptidase M20 domain-containing protein 2 Peptidase M20 domain-containing protein 2
Endopeptidase La Endopeptidase La
Site-2 protease family protein Site-2 protease family protein
Peptidase family M50 Peptidase family M50
Peptidase-M3 domain-containing protein Peptidase-M3 domain-containing protein
L,D-transpeptidase L,D-transpeptidase
Peptidase-A24 domain-containing protein Peptidase-A24 domain-containing protein
lspA Lipoprotein signal peptidase
Extracellular
peptidase S8 peptidase S8
group_294 peptidase S8
group_1522 peptidase S8
group_2124 Peptidase S8
group_2125 peptidase S8
Peptidase S8 Peptidase S8
aprE peptidase S8
acm serine protease like S8
group_5120 peptidase S8
Subtilisin peptide S8
Peptidase-S8 domain-containing protein Peptidase-S8 domain-containing protein
eMpr V8-like Glu-specific endopeptidase
Minor extracellular protease vpr Minor extracellular protease vpr
Extracellular basic protease Extracellular basic protease
Both/Unknown
Di- and tripeptidase Di- and tripeptidase
prolyl aminopeptidase prolyl aminopeptidase
serine protease serine protease
Peptidase S1 domain-containing protein Peptidase S1 domain-containing protein
Peptidase-M78 domain-containing protein Peptidase-M78 domain-containing protein
group_2044 peptidase
group_2073 peptidase
zinc metalloprotease zinc metalloprotease
Peptidase C51 domain-containing protein Peptidase C51 domain-containing protein
Peptidase Peptidase
group_2882 Peptidase
group_2894 peptidase
group_2945 peptidase
peptidase peptidase
group_2952 peptidase
group_2963 peptidase
group_2964 peptidase
Zn-dependent proteases Zn-dependent proteases
group_5239 zinc metalloprotease
group_6644 serine protease
X-Pro dipeptidyl-peptidase (S15 family) domain protein X-Pro dipeptidyl-peptidase (S15 family) domain protein
Peptidase-M23 domain-containing protein Peptidase-M23 domain-containing protein
Peptidase C56 Peptidase C56
Peptidase-S9 domain-containing protein Peptidase-S9 domain-containing protein
Pyroglutamyl-peptidase I Pyroglutamyl-peptidase I
Zinc-dependent metalloprotease Zinc-dependent metalloprotease
Ara h 1
Ara h 2
Gene presence
No
Yes
Degradation scale
Min
Max
Allergen degradation capacity
(western blotting)

**Figure S7.** Protease repertoire of *Rothia* species identified by whole-genome sequencing. Heatmap showing predicted proteases encoded in *Rothia* genomes, including *Rothia mucilaginosa* (R3), *Rothia dentocariosa* (R2), and *Rothia aeria* (R1), as well as additional species without specific isolate names. Rows represent protease genes grouped by subcellular localization (intracellular, extracellular, or both/unknown) with functional annotations, and columns correspond to individual *Rothia* genomes. Blue shading indicates gene presence (light blue: not present, dark blue: present). Green-highlighted labels denote duplicate recognition of the same protease across different sequence annotations. Allergen degrading capacity against Ara h 1 and Ara h 2, determined by Western blotting, is shown at the bottom right for reference and the heat map colour scale (blue-red) represents the percentage of allergen degradation relative to control non-growth incubations (crude peanut extract).

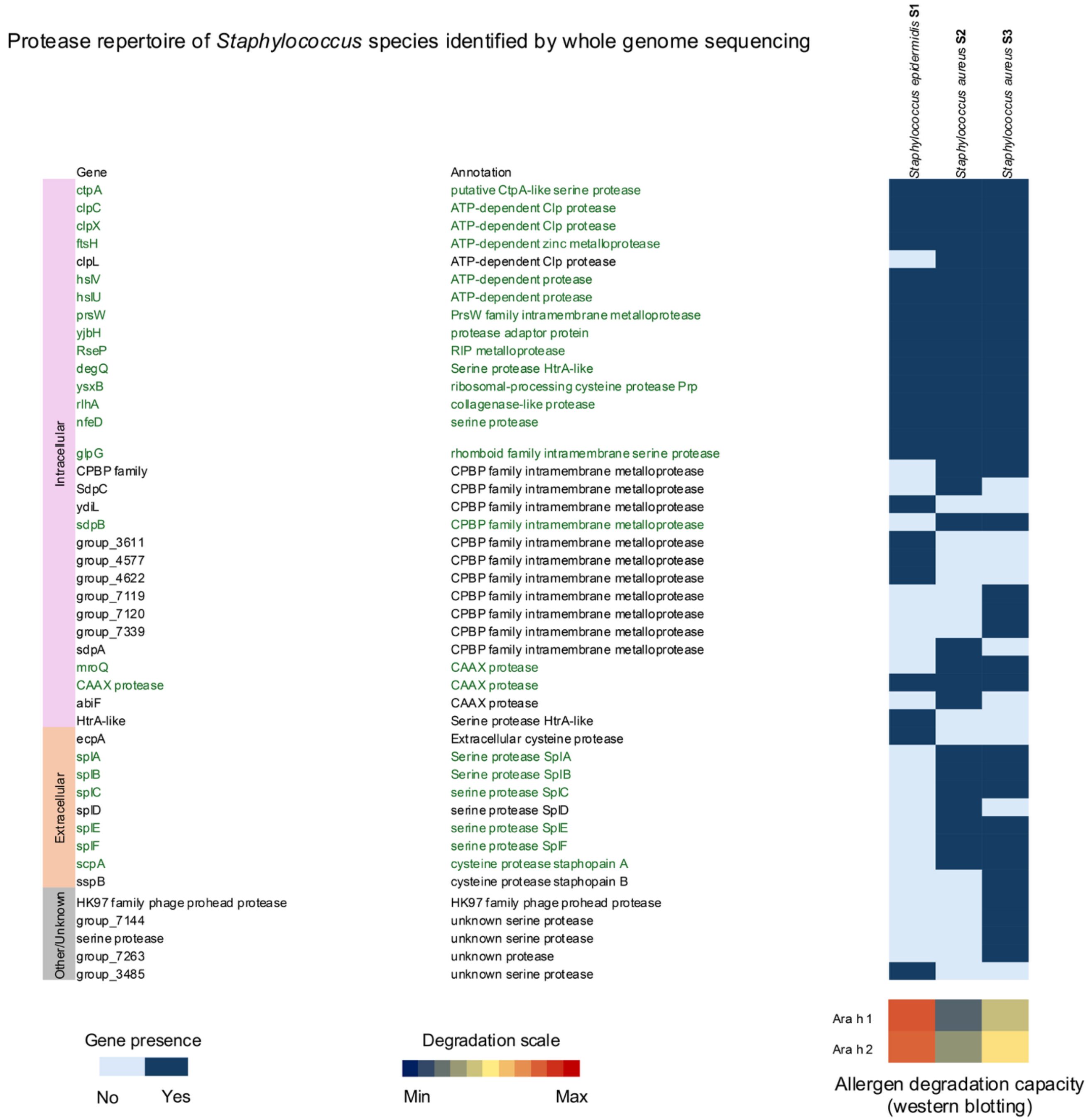


**Figure S8.** Protease repertoire of *Staphylococcus* species identified by whole-genome sequencing. Heatmap showing predicted proteases encoded in *Rothia* genomes, including *Staphylococcus epidermidis* (S1), *Staphylococcus aureus* (S2), and *Staphylococcus aureus* (S3). Rows represent protease genes grouped by subcellular localization (intracellular, extracellular, or both/unknown) with functional annotations, and columns correspond to individual *Staphylococcus* genomes. Blue shading indicates gene presence (light blue: not present, dark blue: present). Green-highlighted labels denote duplicate recognition of the same protease across different sequence annotations. Allergen degrading capacity against Ara h 1 and Ara h 2, determined by Western blotting, is shown at the bottom right for reference and the heat map colour scale (blue=0%, red=100%) represents the percentage of allergen degradation relative to control non-growth incubations (crude peanut extract).

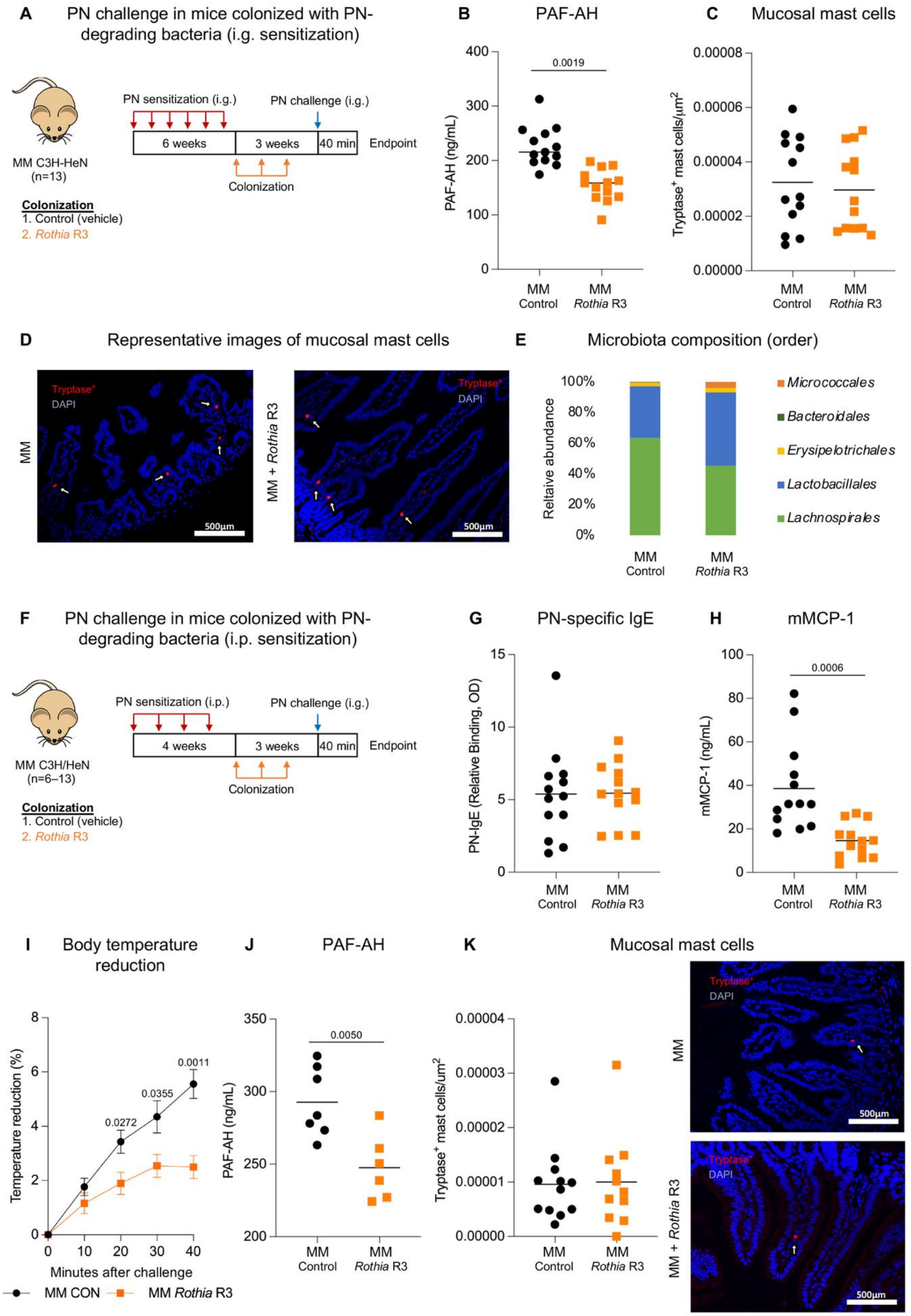

A
PN challenge in mice colonized with PN-degrading bacteria (i.g. sensitization)
PN sensitization (i.g.)
PN challenge (i.g.)
6 weeks
3 weeks
40 min
Endpoint
Colonization
MM C3H-HeN
(n=13)
Colonization
1. Control (vehicle)
2. Rothia R3
B
PAF-AH
PAF-AH (ng/mL)
0.0019
MM Control
MM Rothia R3
C
Mucosal mast cells
Tryptase+ mast cells/µm2
MM Control
MM Rothia R3
D
Representative images of mucosal mast cells
Tryptase+
DAPI
MM
MM + Rothia R3
500µm
E
Microbiota composition (order)
Reltaive abundance
Micrococcales
Bacteroidales
Erysipelotrichales
Lactobacillales
Lachnospirales
MM Control
MM Rothia R3
F
PN challenge in mice colonized with PN-degrading bacteria (i.p. sensitization)
PN sensitization (i.p.)
PN challenge (i.g.)
4 weeks
3 weeks
40 min
Endpoint
Colonization
MM C3H/HeN
(n=6–13)
Colonization
1. Control (vehicle)
2. Rothia R3
G
PN-specific IgE
PN-IgE (Relative Binding, OD)
MM Control
MM Rothia R3
H
mMCP-1
mMCP-1 (ng/mL)
0.0006
MM Control
MM Rothia R3
I
Body temperature reduction
Temperature reduction (%)
Minutes after challenge
0.0272
0.0355
0.0011
MM CON
MM Rothia R3
J
PAF-AH
PAF-AH (ng/mL)
0.0050
MM Control
MM Rothia R3
K
Mucosal mast cells
Tryptase+ mast cells/µm2
MM Control
MM Rothia R3
Tryptase+
DAPI
MM
MM + Rothia R3
500µm

**Figure S9.** (A–D) Peanut (PN) intragastric (i.g.) sensitization and challenge in minimal microbiota (MM) mice colonized with or without *Rothia* R3, continued from Figure 7. (A) Experimental design. MM C3H/HeN mice were sensitized i.g. to PN weekly for 6 weeks, then colonized with *Rothia* R3 weekly for 3 weeks. PN-challenge was given i.g. 3 weeks after the first colonization, followed by sacrifice 40 min later. n=13 mice/group; pooled from 3 independent experiments. (B) Serum platelet activating factor acetylhydrolase (PAF-AH). (C) Tryptase-positive mast cells quantified in small intestinal sections. (D) Representative images of small intestinal mucosal tryptase-positive mast cells. (E) Small intestinal microbiota composition (order). (F–H) PN i.p. sensitization and i.g. challenge in MM mice colonized with or without *Rothia* strains. (F) Experimental design. MM C3H/HeN mice were sensitized i.p. to PN weekly for 4 weeks, then colonized with *Rothia* R3 weekly for 3 weeks. PN-challenge was given i.g. 3 weeks after the first colonization, followed by sacrifice 40 min later. (G) Serum PN-specific IgE before PN challenge. (H) Serum mucosal mast cell protease 1 (mMCP-1) after PN challenge. (I) Core body temperature reduction after PN challenge. (J) Serum PAF-AH after PN challenge. (K) Tryptase-positive mast cells quantified in small intestinal sections with representative images. n=13 mice/group; pooled from 3 independent experiments (G, H, K). n=6 mice/group pooled from two independent experiments (I–J). Displayed *P* values were calculated using an unpaired Student's *t*-test.

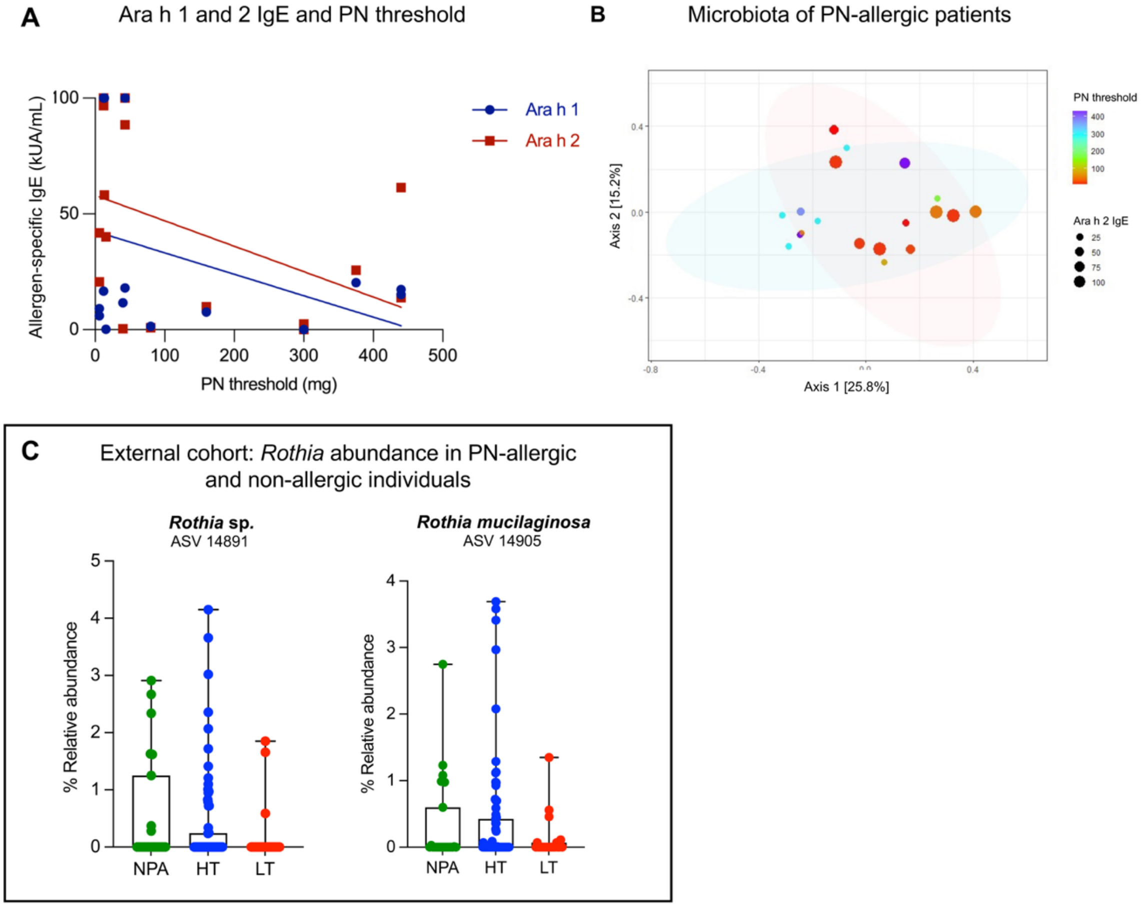


**Figure S10.** (A–B) Peanut (PN)-allergic patients (n=19) were tested for PN threshold using increasing PN concentrations before enrolment in oral immunotherapy. (A) Correlation plot of Ara h 1-specific IgE ($y$= -0.09$x$ + 42.45, $R^2$ = 0.15, $P$=0.11) and Ara h 2-specific IgE ($y$= -0.11$x$ + 58, $R^2$= 0.21, $P$=0.06) against PN threshold. (B) Bray-Curtis-tom represents serum Ara h 2-specific IgE levels, while dot color corresponds to the PN threshold (scales shown). (C) Abundance of *Rothia* species in saliva from analysis of 16s rRNA data from an external cohort including 120 children (23 non-PN-allergic controls (NPA), 74 PN-allergic with high PN threshold (HT; >443 mg), and 23 PN-allergic with low PN threshold (LT; <443 mg)) who underwent double-blind, placebo-controlled PN challenges. Data are presented as interquartile range (IQR) with whiskers extending to the min and max data points.

**Table S1: Detailed subject demographics. Peanut, PN; Male, M; Female; F.**

| Status | ID | Sex | Age | Race | PN threshold (mg) | Ara h 1 IgE (kUA/mL) | Ara h 2 IgE (kUA/mL) |
|---|---|---|---|---|---|---|---|
| PN-allergic saliva | PN1 | F | 10 | Unknown | 300 | 0.10 | 1.19 |
| | PN2 | F | 7 | Asian | 300 | 0.10 | 2.44 |
| | PN3 | F | 1 | White | 300 | 0.10 | 0.10 |
| | PN4 | F | 27 | White | 440 | 15.2 | 13.8 |
| | PN5 | F | 6 | White | 375 | 20.3 | 25.7 |
| | PN6 | M | 6 | White | 40 | 11.6 | 0.41 |
| | PN7 | M | 7 | White | 300 | N/A | N/A |
| | PN8 | F | 15 | White | 440 | 17.3 | 12.1 |
| | PN9 | M | 9 | White | 12 | 100 | 25.8 |
| | PN10 | M | 13 | White | 6 | 9.11 | 0.19 |
| | PN11 | M | 10 | White | 43 | 18 | 88.5 |
| | PN12 | F | 13 | White | 13 | 100 | 58.2 |
| | PN13 | F | 13 | White | 43 | 100 | 100 |
| | PN14 | F | 14 | White | 12 | 100 | 96.7 |
| | PN15 | M | 9 | White | 6 | 5.95 | 20.7 |
| | PN16 | M | 11 | Asian | 160 | 7.62 | 9.92 |
| | PN17 | F | 2 | Asian | 80 | 1.41 | 0.84 |
| | PN18 | F | 2 | Black/African American | 15.5 | 0.18 | 40.1 |
| | PN19 | M | 4 | Unknown | 12 | 16.7 | 100 |
| Non-PN-allergic saliva | HV1 | M | 35 | White | N/A | N/A | N/A |
| | HV2 | M | 22 | White | | | |
| | HV3 | M | 20 | White | | | |
| | HV4 | F | 26 | White | | | |
| | HV5 | F | 33 | White | | | |
| | HV6 | F | 24 | White | | | |
| | HV7 | F | 18 | White | | | |
| | HV8 | M | 30 | White | | | |
| | HV9 | F | 38 | White | | | |
| | HV10 | F | 20 | Black/African American | | | |
| | HV11 | M | 40 | Asian | | | |
| | HV12 | F | 66 | White | | | |
| | HV13 | M | 33 | White | | | |
| Non-PN-allergic jejunal washes | HV14 | M | 57 | White | N/A | N/A | N/A |
| | HV15 | M | 63 | White | | | |
| | HV16 | F | 57 | White | | | |
| | HV17 | F | 35 | White | | | |
| | HV18 | M | 53 | White | | | |

**Table S2: Serum characterization for western blotting. Peanut, PN.**

| Serum | Total IgE levels (kU/L) | PN IgE (kUA/mL) | rAra h 1 IgE (kUA/mL) | rAra h 2 IgE (kUA/mL) | rAra h 3 IgE (kUA/mL) | rAra h 8 IgE (kUA/mL) | rAra h 9 IgE (kUA/mL) | Sex | Age | Positive Skin Prick Test | Other hypersensitivities |
|---|---|---|---|---|---|---|---|---|---|---|---|
| **A** | 12.49 | Unknown | 52.9 | >100 | 26.6 | 0.02 | 0.02 | F | 30 | PN, pollen of Cupressaceae, Acer pseudoplatanus, olive, grasses, Salsola, dog, cat, horse and rabbit. Minimal reaction to mites and aspergillus | Persistent moderate asthma and intermittent rhinoconjunctivitis |
| **B** | 176 | 54.9 | 20.4 | 24.5 | 0 | 0 | 0 | F | 19 | PN, chickpea, arizonica pollen, minimal reaction to pine nuts | Atopic dermatitis |
